# The statistics of noisy growth with mechanical feedback in elastic tissues

**DOI:** 10.1101/426361

**Authors:** Ojan Khatib Damavandi, David K. Lubensky

**Affiliations:** Department of Physics, University of Michigan, Ann Arbor MI, USA 48109-1040

**Keywords:** tissue mechanics, noisy growth, clonal dynamics, mechanical feedback

## Abstract

Tissue growth is a fundamental aspect of development and is intrinsically noisy. Stochasticity has important implications for morphogenesis, precise control of organ size, and regulation of tissue composition and heterogeneity. Yet, the basic statistical properties of growing tissues, particularly when growth induces mechanical stresses that can in turn affect growth rates, have received little attention. Here, we study the noisy growth of elastic sheets subject to mechanical feedback. Considering both isotropic and anisotropic growth, we find that the density-density correlation function shows power law scaling. We also consider the dynamics of marked, neutral clones of cells. We find that the areas (but not the shapes) of two clones are always statistically independent, even when they are adjacent. For anisotropic growth, we show that clone size variance scales like the average area squared and that the mode amplitudes characterizing clone shape show a slow 1*/n* decay, where *n* is the mode index. This is in stark contrast to the isotropic case, where relative variations in clone size and shape vanish at long times. The high variability in clone statistics observed in anisotropic growth is due to the presence of two soft modes—growth modes that generate no stress. Our results lay the groundwork for more in-depth explorations of the properties of noisy tissue growth in specific biological contexts.

---

Growth is central to biology and usually involves a level of stochasticity (1–3). The presence of such noise can have significant consequences for developmental processes like morphogenesis and the regulation of organ size. Yet, relatively little is known about quantitative aspects of stochastic growth. Our goal here is to understand the interplay between noise and mechanical feedback in growing elastic tissues. To that end, we present a generic, continuum model of such tissues and use it to study measurable features of tissue architecture like density fluctuations and statistics of marked, neutral clones. Our model makes a number of unexpected predictions, including the presence of power law correlations in space and the existence of soft modes that allow clone sizes to grow diffusively, evading the effects of mechanical feedbacks that might otherwise be expected to limit clone size variability.

In experiments, noise in growth has most often been probed through the size and spatial distribution of clones of cells (4–10), especially of neutral clones that are genetically identical to surrounding tissue except for a clone marker. Cell density variation has also been observed directly in culture (11–15) and in fixed tissues (16), as have correlations in positions of mitotic cells (1), and size asymmetry between contralateral organs can be used as an indirect readout of noise levels (17).

Theoretically, the most thoroughly explored area is the noisy dynamics of stem cell populations (4, 5, 9), where zero-dimensional descriptions are often appropriate and spatial crowding effects, when important, have been included at the level of simple lattice models. Similar models apply to fluid tissues where clone fragmentation and aggregation is the dominant process at long times (6). Here, in contrast, our goal is to explicitly include the effects of mechanical stresses on growth of elastic tissues. An elastic description is typically valid for plants (18, 19) and can be applied to animal tissues like the *Drosophila* wing imaginal disc to the extent that cell rearrangements are rare (20–24); moreover, the formalism we adopt can be extended to encompass simple models of plastic flow that allow for more frequent rearrangements (25). We expect that noise will lead to growth non-uniformities and thus to the uneven accumulation of cell mass; this, in turn, will generate stresses that can feed back on local growth rates. Ranft *et al*. (26) have shown that such mechanical feedbacks cause the stress tensor to relax as if the tissue were a viscoelastic fluid; their treatment of noise, however, is limited to fluctuations about zero average growth, where most of the phenomena of interest here are absent.

On a very basic level, the idea of mechanical feedback on growth is uncontroversial—cells obviously cannot grow indefinitely into space occupied by other cells, so some sort of contact inhibition or crowding effects must be present. Whether cells more generally adjust their growth rates to their mechanical environment is, however, less obvious. The idea that negative feedback from mechanical stresses could damp out cell density fluctuations was proposed in (27) and has since been incorporated into a number of models (8, 18, 19, 28–37). Clear experimental examples of mechanical feedback have been more difficult to obtain (38, 39). Nonetheless, several studies have argued that tissues both in culture (12, 40–42) and *in vivo* (43–47) respond to mechanical stress by modulating the rate and orientation of cell division or by inducing cell death (42, 48–50). Clones of fast-growing cells in *Drosophila* wing discs reduce their growth rate through mechanical feedback (8), and similar behavior has been observed in plant systems (19, 51) including the *Arabidopsis* sepal (10). In confluent monolayers, contact inhibition slows mitosis (11, 15). Cell aggregates (52–55) and bacterial populations (56) also appear to respond to mechanical cues. Thus, it seems likely that some mechanical feedback on growth is present in many tissues.

In what follows, we first introduce our basic framework, which assumes linear elastic deformations about a uniformly growing reference state and linear feedback of the stress tensor on the local growth rate. We then consider the special case of strictly isotropic growth, where we show that density-density correlations generically decay with distance as a power law and that mechanical feedback drives clone size variability to zero on large scales. (Closely related results are obtained independently in (57).) We also find that the areas of neighboring clones are statistically uncorrelated. We then turn to the more general case of locally anisotropic growth. Here, we observe the appearance of soft modes, where faster growth of one tissue region is exactly compensated by slower growth in surrounding regions and elastic deformations so as to leave the tissue completely stress-free; these modes can thus grow without bound even in the presence of feedback. As a consequence, clone sizes display a standard deviation of order their mean, in strong contrast to the isotropic case.

## Basic Model

We consider a flat epithelium undergoing isotropic, exponential growth on average, with small, random deviations from this average. Though many of our results can be generalized, we limit ourselves here to two-dimensional tissues. At the macroscopic scale, we view the epithelium as an elastic continuum. Growth then looks like local creation of mass, and non-uniform growth can induce tissue deformations. We treat the simplest case of an infinite tissue and ignore boundary effects.

With these assumptions, we define reference (Lagrangian) coordinates **{R}** and material (Eulerian) coordinates γ. Each point on the tissue at time *t* is related to a point in the reference coordinates by 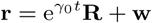, where *γ*_0_ is the average growth rate and **w** the local deformation due to growth fluctuations (Fig. 1). We focus primarily on *γ* _0_ *>* 0 but show in the Supporting Information (SI) that we recover known results (26) when *γ* _0_ = 0.

**Fig. 1.**
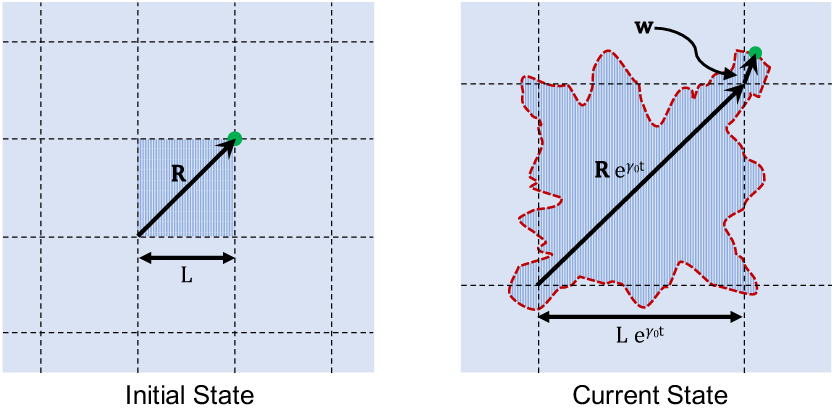
Noisy growth leads to tissue deformation. A shaded square of side *L* in the initial state has grown at time *t* into a larger region with deformed boundaries (red); dashed grid lines indicate the average, uniform tissue dilation. A material point (green dot) initially at **R** is displaced to 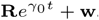.

Growth is represented by the symmetric second rank tensor G(**R**) (see, e.g., (25)). Its principal components give the tissue’s preferred dilation—that is, the factor by which the tissue particle at **R** must expand to remain stress free—along the two principal axes. It can be decomposed as 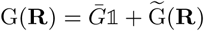, where the scalar 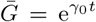 describes the spatially uniform, average tissue expansion, and the tensor 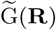 represents fluctuations about the average.

We assume that growth is slow enough that the tissue is always at mechanical equilibrium; absent any external forces or confinement, spatially uniform growth then should not generate any mechanical stress, which will instead be caused entirely by the spatially-varying component 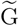 of the growth tensor. In the limit of weak fluctuations in growth rate, 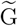 and **w** will both be small, and we can linearize at each instant about the uniformly dilated state. The resulting theory has a form familiar from thermoelasticity (58, 59). We define a strain-like tensor *w*_*ij*_ = (*∂*_*i*_*w*_j_ + *∂*_*j*_ *w*_*i*_)*/*2, where *∂*_*i*_ denotes the partial derivative with respect to *R*_*i*_. (Throughout this paper, spatial derivatives are taken with respect to Lagrangian coordinates unless otherwise specified.) The Cauchy stress tensor is then:

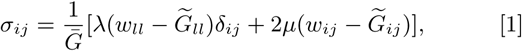

where *λ* and *µ* are the Lamé coefficients and summation on repeated indices is implied. Intuitively, Eq. 1 says that the stress vanishes when the actual strain *w*_*ij*_ matches the preferred local deformation due to growth 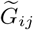 and otherwise grows linearly with the difference between these two quantities. It differs from textbook thermoelastic results only in the factor of 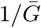, which arises from the conversion of derivatives with respect to **R** to derivatives with respect to the uniformly dilated reference coordinates 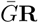 Eq. 1 can also be obtained by linearizing a fully nonlinear theory of morphoelasticity (25) (see SI).

Given Eq. 1, force balance *∂iσ*_*ij*_ = 0 implies

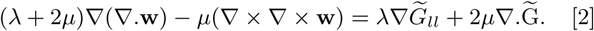

This equation may be solved to obtain **w**, and thus *w*_*ij*_ and, *σ*_*ij*_ at each instant as a function of 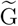.

To complete our description, we must specify the dynamics of G, whose most general possible form is

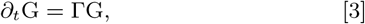

where Γ is a rank four tensor that incorporates mechanical feedback and noise (see SI). We define *∂*_*t*_ to be a time derivative taken at fixed Lagrangian coordinates **R**. (The SI discusses how *∂*_*t*_G with this convention is related to proposed expressions for the time derivative at fixed Eulerian coordinates.)

Assuming that deviations from uniform growth are small, we can expand Eq. 3 to linear order in stress feedbacks and noise. The most general form allowed by symmetry is

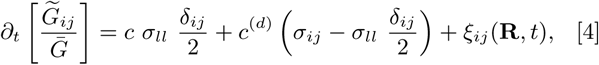

where we have also dropped higher order gradient terms (whose effects are addressed in the SI). The constants *c* and *c*^(*d*)^ give the strengths of the stress feedbacks (the superscript (*d*) stands for deviatoric), and *ξ*_*ij*_ is a noise term.

### Density

Within continuum elasticity, it is natural to define a density *ρ*(**R**, *t*) of the deformed material. For a complex biological tissue, *ρ* of course does not represent the total mass density but instead can be thought of as roughly the density of materials (like cytoskeletal proteins) that give the tissue an elastic rigidity or, equivalently, as a measure of elastic tissue compression or expansion relative to an ideal, unstressed state. If we call the (uniform) density in the stress-free configuration *ρ*_0_ and deviations *δρ*(**R**, *t*) = *ρ*(**R**, *t*) *-ρ*_0_, then to linear order in growth fluctuations,

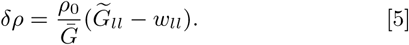

If all cells in the tissue have the same preferred apical area, then *ρ* is proportional to the cell density. Even if that is not the case, as long as cells’ relaxed areas are uncorrelated, the difference between the cell density and t<u>h</u>e average of *ρ* over a region of area *A* will decrease as 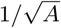, allowing the cell density to be used to estimate *ρ* on long enough scales.

## Isotropic Growth

In order to build intuition, we first consider the simplest case of isotropic growth. We thus set 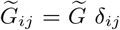 and keep only the isotropic part of Eq. 4 by taking *c*^(*d*)^ = 0 and *ξ*_*ij*_ = *ξδ*_*ij*_ */*2. As outlined above, we can use force balance (Eq. 2) to find *w*_*ij*_ in terms of 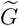. After Fourier transforming, we have 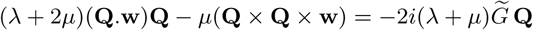. Solving for **w** and using Eq. 5 to relate *δρ* and 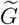, we find

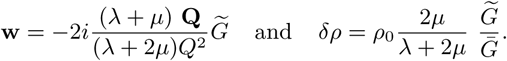

Then Eq. 4 can be rewritten as

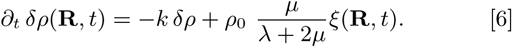

Here, *k* = 2*µ*(*λ* + *µ*)*c/*(*λ* + 2*µ*), and the (scalar) noise ***ξ***(**R**, *t*) is delta correlated in time and spatially correlated on a length scale *a* of order a cell radius; this small but non-zero correlation length is needed to prevent pathological behavior and reflects the fact that cell growth and division are correlated on the scale of a single cell. Because typical cell sizes remain constant as the tissue grows, the correlation length should be viewed as fixed in Eulerian coordinates, so that 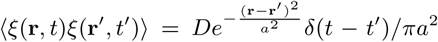, where *D* is the noise strength. In Lagrangian coordinates, the noise then satisfies 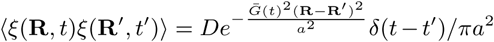, where we used, to leading order, 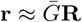.

### Density-density Correlations

From Eq. 6 we can calculate the density-density correlation function, which we find in steady state drops off like a power law in *r* for large separations *r ≫ a:*

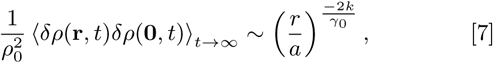

where the pre-factor is *𝒪* (*k/γ*_0_) (see SI). Fig. S1 plots the full correlation function versus *r/a*.

To understand this behavior, suppose that noise generates a density fluctuation initially correlated on a length scale *a*. Over time, growth will advect this fluctuation outwards while mechanical feedback will cause its amplitude to decay. After time *t*, the initial, small-scale fluctuation will induce correlations up to a scale 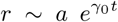, but *δρ* will have decayed like *e*^*-kt*^ (so that the correlation function is smaller by a factor *e*^*-*^2^*kt*^). For a given *r*, fluctuations that happened around *t*^*∗*^(*r*) = log(*r/a*)^1*/γ*^_0_ ago are dominant; earlier fluctuations have died out while later ones have not yet reached the distance *r*. Thus, we expect 〈*δρ*(**r**)*δρ*(0) 〉 *∼* exp[*-*2*kt*^*∗*^(*r*)] *∼* (*r/a*)^*-*2*k/γ*^_0_. A similar mechanism produces power law correlations in inflationary models of the early universe (60). This result requires a small but finite initial correlation length for fluctuations. In the SI, we extend our results to the case where this lengthscale is set by gradient terms in the growth dynamics as well as to unequal time correlations in arbitrary dimension.

Qualitatively, our finding of power law correlations says that cells in growing epithelia are more clumped together on large scales than for a totally random spatial distribution, with the effect becoming more pronounced for smaller *k/γ*0. Correlations in cell density are experimentally accessible (13– 16), and we expect that similar clumping should be observed in spatial distributions of mitotic cells (which, intriguingly, are known cluster in some tissues (1, 24)). Estimates of the exponent 2*k/γ*_0_ could be used to determine the strength *c* of mechanical feedback, which has not previously been measured.

### Clone Statistics

Following marked, neutral clones has proven to be a useful tool to track growth and development (4–10). Here, we examine the size and shape statistics of clones in a growing tissue. Starting from a circular clone, we derive expressions for the variance of the clone area and of mode amplitudes characterizing the clone shape.

The area of a clone with initial radius *R*_*c*_ is 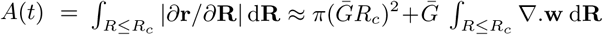, to leading order in small *w*. The variance of clone size is then

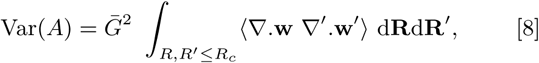

where *Δ’* = *Δ*_R’_ is the gradient operator taken with respect to **R**’. To study clone shape, we describe the instantaneous clone boundary as a curve *r*_*c*_(*θ*) in Eulerian coordinates, where *θ* is the polar angle, which we can then express as a Fourier series *r*_*c*_(*θ*) = ∑*B*_*n*_*e*^*inθ*^. To linear order in **w**, we may neglect differences between the Lagrangian and Eulerian polar angles Θ and *θ*, and 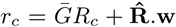, where 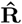 is a unit vector in the direction of **R**. Continuing to work to the same order, and defining 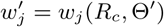, it is easy to see that

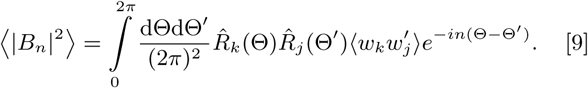

For isotropic growth, both **w** and ∇.**w** can be written in terms of *δρ*, so Eq. 6 is all we need to evaluate Eqs. 8 and 9. We find that the variances in clone size and shape become small compared to the mean at long times: compared to the meanat long times: lim_t → 221E;_ Var(***A***)/〈***A***〉^2^ → 0 and 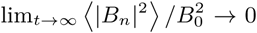. Thus, for isotropic growth, all sufficiently old clones are statistically similar. This is a direct consequence of the mechanical feedback that causes *δρ* to relax exponentially to zero; we will see in the next section that once growth is allowed to be locally anisotropic, soft modes lead to dramatically different behavior.

Before introducing anisotropy, we note that our model also predicts that the areas of two nearby clones are uncorrelated: If the deviations from the mean clone areas are Δ*A*_1_ and Δ*A*_2_, then 〈 Δ ***A*_1_** Δ ***A*_1_ 〉 =0.** This follows immediately from the elastic response of an infinite tissue to localized growth (8): With 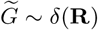, 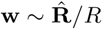 and *∇.***w** = 0 except at the origin. Thus, while the shapes of clones near the origin are distorted, only clones that actually contain the origin change their area (Fig. 2A). More generally, for both isotropic and anisotropic growth, the size of each clone depends only on 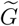 within its own boundaries, and 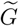 is spatially uncorrelated (see SI).

**Fig. 2.**
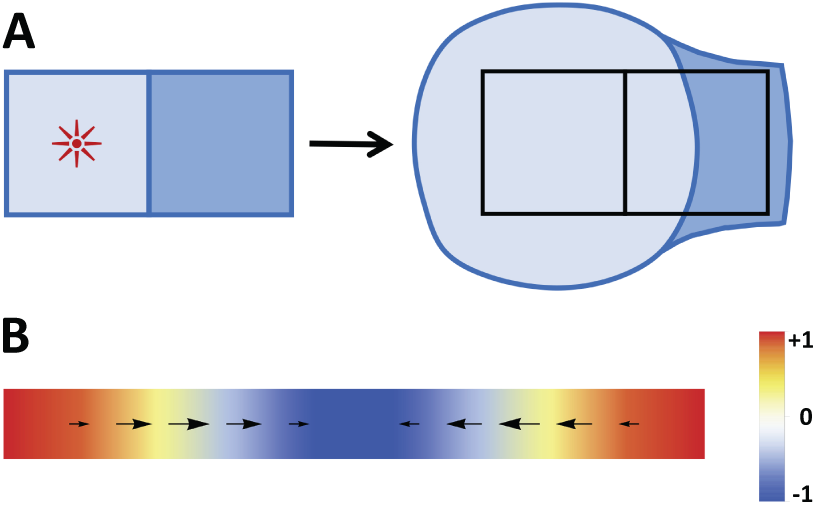
(A) Independence of nearby clone areas. A localized region of growth (marked by the red star) at the center of the left clone (light blue) leads to an increase in the area of that clone, but it leaves the area of the adjacent clone (dark blue) unchanged even while distorting its shape. (B) Longitudinal soft mode. Sinusoidal growth (color scale gives 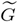) leads to a deformation field **w (**arrows) that exactly compensates for the growth, leaving the density unchanged.

## Anisotropic Growth

Cells in both animal (41, 43, 44) and plant (51) tissues have been shown to orient their divisions relative to the principal axis of an applied stress. Thus, in general, we should consider stress-dependent, local growth anisotropies. If we continue to assume that growth is isotropic on average so that 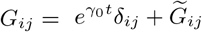, then we can specify the symmetric tensor 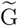 by three scalars: its trace and two components for the traceless part. We follow the same steps as in the isotropic case to solve Eq. 4 and find expressions for density-density correlations and clone statistics. We will see that density-density correlations follow the same power law behavior as in Eq. 7, but that new soft modes appear which cause clone size and shape variability to remain large even at long times.

To solve Eq. 2 for **w** in terms of 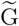, we work in Fourier space and write the traceless part of 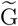 in terms of the *Q*-dependent scalars 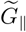 and 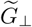 (see also the SI):

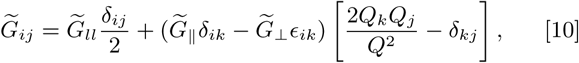

where ∈_*ij*_ is the Levi-Civita tensor. Eq. 2 then gives the components of **w** longitudinal and transverse to **Q**:

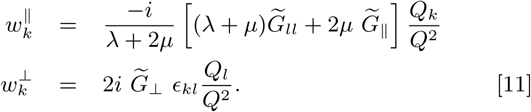

Finally, we have the stress tensor:

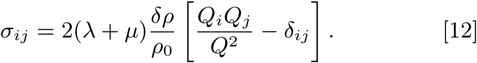

Strikingly, growth in an infinite system can thus only induce a nonzero stress when there is a non-vanishing density fluctuation *δ*_*ρ*_ related to 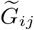 by

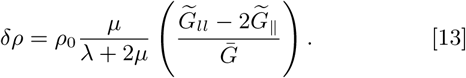

Having found the stress, we next turn to the time evolution of G. The noise *ξ*_*ij*_ in Eq. 4 now has three independent components. We define *ξ*_*ll*_, *ξ* _*‖*_ and *ξ*_*⊥*_ in analogy to the corresponding quantities in Eq. 10 and take them to be independent, Gaussian random variables; rotational invariance requires that *ξ* _*‖*_ and *ξ*_*⊥*_ have the same strength which can, however, differ from that of *ξ*_*ll*_. As in the isotropic case, we choose these random variables to be delta correlated in time but colored in space to avoid pathological behavior (see SI).

Substituting Eq. 12 for the stress into the growth dynamics Eq. 4, we find that the dynamics can be decomposed into three independent modes as (see also SI)

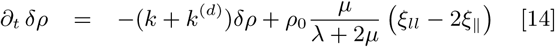

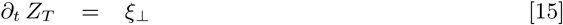

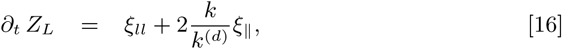

where *k*^(*d*)^ is defined similarly to *k*, and 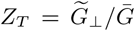 and 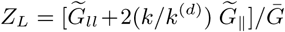 are the amplitudes respectively of transverse and longitudinal soft modes, which do not produce any stress and thus grow diffusively. Before studying the density and clone statistics, we next consider these soft modes more carefully.

### Soft Modes

As just shown, in two dimensions growth has two soft modes. They have the same physical origin: A nonuniform growth field 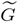 induces a displacement **w** that exactly cancels the growth, so that no mechanical stress or density change results. Fig. 2B illustrates this for the longitudinal mode. In essence, excess mass created by faster growth in one part of the tissue is transported to slower-growing parts so as to equalize the tissue density. What is remarkable is that, for certain patterns of growth, this redistribution can be accomplished in an elastic tissue—thus without viscous flow or cell neighbor exchanges—without causing shear stresses or mechanical feedback. Specifically, Eq. 13 implies that only growth perpendicular to **Q**—i.e., only the 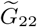 component of 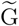, if **Q** points in the 1 direction—induces density changes that can then generate stresses (Eq. 12). In contrast, growth parallel to **Q (***Z*_*L*_) or at a 45° angle (*Z*_*T*_) is exactly compensated by elastic deformations.

From Eqs. 15–16, one might imagine that the soft modes grow without bound. This, however, turns out not to be the case: The modes are defined at fixed Lagrangian wavevector **Q**, but *ξ*_*⊥*_ and *ξ* _‖_ have constant correlations in Eulerian space. As growth progresses, a given **Q** corresponds to longer and longer Eulerian lengthscales, leading to a decrease in the noise amplitude with time. As a result (see SI), the mean squared values of *Z*_*L*_ and *Z*_*T*_ remain bounded for all times.

### Density-density Correlations

As in the isotropic case, we can calculate density-density correlation functions for anisotropic growth. From Eq. 14, it is evident that the only differences between the two cases are the feedback strength and the presence of a second noise term *ξ* _*‖*_ The latter will only affect the pre-factors. We thus find that

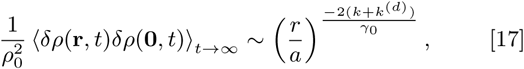

just as for isotropic growth, with prefactors of the same order.

### Clone Statistics

In the isotropic case, where *δ*_*ρ*_ was the only dynamical variable, mechanical feedback caused clone size and shape variations to decay in time relative to the average dilation. However, if we allow growth anisotropies, ∇.**w** and **w**, which are the relevant variables for clone properties (Eqs. 8 and 9), depend on the soft modes as well as on the density.

As mentioned above, the correlation functions for soft modes reach constant values at long times, whereas correlators that involve *δ*_*ρ*_ decay exponentially in time. Thus ⟨∇.**w** ∇′.**w**′⟩ and 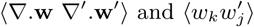 will be dominated by soft modes at long times. We find that as *t* → ∞ the clone area obeys

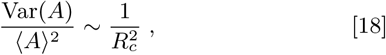

while for *n* ≥ 2 the mode amplitudes show a slow 1*/n* decay:

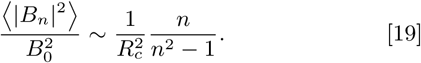

The pre-factors are *𝒪*(1) if *R*_*c*_ ∼ *a*, i.e. the clone is initially the size of a single cell. Adjacent clones’ areas are uncorrelated, just as in the isotropic case (Fig. 2A).

Eq. 18 says that the areas of large clones are just as variable as those of small clones. Such behavior would be normal for an exponentially growing population with no constraints on its size (61), but it is quite unexpected for a clone embedded in an elastic tissue and subject to mechanical feedback. In effect, soft modes allow the growth to be as noisy as if feedback were absent. Together with our finding that the areas of different clones are uncorrelated, this result means that even elastically coupled clones behave in many ways as if they were growing independently and without feedback.

## Discussion

We presented a simple model of noisy growth of elastic tissues with mechanical feedback. This model predicts nontrivial behavior for experimentally accessible quantities including density-density correlation functions and clone size statistics.

We first showed that density correlations decay in space as a power law whose exponent depends on the average growth rate and feedback coefficient. Using our result, it should be possible to estimate the strength *c* + *c*^(*d*)^ of the mechanical feedback, a quantity that has not so far been measured in growing tissues. Somewhat counterintuitively, the sign of *c*^(*d*)^ seems to be negative in some plant systems (51). We find that this does not destabilize growth as long as *c* + *c*^(*d*)^ > 0.

We then studied marked, neutral clones, whose statistics depend strongly on whether local growth anisotropies are allowed. In their absence, mechanical feedback tends to prevent fluctuations on large scales, and size variation becomes much smaller than the mean for large clones. This behavior changes dramatically for anisotropic growth, where soft modes allow certain deformations to escape any negative feedback. As a consequence, the standard deviation in clone area grows like the mean, exactly as if the clone were an exponentially growing aggregate of independently dividing cells completely indifferent to the presence of surrounding tissue (61). Similarly, fluctuations in clone shape remain large when growth is anisotropic and soft modes dominate. Strikingly, we also find that the areas of different clones are always uncorrelated. These conclusions together imply that (at least to within the weak noise approximation inherent in our calculations) neutral clone areas have the same statistical properties in elastic tissues as in systems where mechanical feedback and crowding effects are completely absent.

Our calculations also show more generally that overgrowth of one clone (whether or not neutral) cannot induce under-growth in nearby tissue solely through mechanical feedback in a linear, elastic continuum (SI and (8)); this is consistent with the fact that known examples of such behavior, like cell competition (62), appear to depend on the activation of specific signaling pathways rather than on generic mechanical effects.

One limitation of our results is that they ignore clone disappearance and fragmentation. Such events should be vanishingly unlikely for fast enough growth or large enough initial clone area, but can skew size distributions in the opposite limits (4). Thus, our findings are most directly applicable to tissues where the rate of cell division is much higher than of cell death or to situations where clones can be imaged over time (7), so that it is possible to quantify a clone’s incremental growth after it has reached some threshold size.

We emphasize that the soft modes in our model are distinct from stress relaxation due to tissue fluidization (26). Whereas in the latter case, stress decays exponentially in time, for soft modes it is identically zero. Our soft modes are related to harmonic growth (28, 63, 64), which likewise does not generate any stress. This phenomenon, however, occurs in finite tissues with isotropic growth and appropriate boundary conditions. Here, anisotropic soft modes are integral to the structure of the growth problem even in the absence of boundary effects.

Our model is the simplest that incorporates noise in the growth control problem. It ignores the influences both of boundaries and of frictional forces. Friction always dominates for large enough growing tissues (65), but its effects can be small when growth is slow or sources of drag are weak (as, e.g., for plant tissues growing in air). By assuming a solid tissue, we also neglect the possibility of cell rearrangements (T1 transitions) and flow. This assumption usually holds for plant cells (18, 19). In animal epithelia, flows and T1’s are sometimes significant (6), but there are also cases that exhibit more solid-like behavior. The *Drosophila* wing disc, for example, clearly supports circumferential stresses without yielding (43, 44), and cell shapes and packing are consistent with a solid rather than a fluid-like phase (66). Despite some differences in measured rates of T1’s in *ex vivo* discs (22, 24) and of clone dispersal (20, 21, 23), these are consistent with predominantly solid behavior with possibly some plastic slippage. Thus, it is reasonable to treat the disc as an elastic solid in a first approximation. Moreover, plastic deformations can be described with the same multiplicative decomposition of the deformation gradient used in morphoelasticity (SI), so that incorporating a simple version of plasticity into our model requires nothing more than renormalizing the coefficients in the time evolution of G (Eq. 4) (25).

Our calculations also assume weak noise and hence small deformations. Importantly, this approximation is valid and self-consistent even with soft modes, because the soft mode amplitude is proportional to the noise strength and remains bounded at long times. Nonetheless, it would be of interest to explore what happens for stronger noise and larger deformations. Indeed, the physics of nonequilibrium growth has a long and rich history (e.g. (67)), and from this perspective volumetric tissue growth represents an entirely new class of problems. Notably, we expect that the basic physics of soft modes survives the transition to the nonlinear regime. In the nonlinear case, it is convenient to characterize growth by a “target metric” (equal to G^*T*^ G); if this metric lacks intrinsic curvature, it can always be compensated by a displacement field **w** and so will not generate any stresses (25, 68).

The formalism presented here can also be extended to include less generic effects, like inhomogeneous growth driven by morphogen gradients and chemical signaling, that are nonetheless crucial to many examples of morphogenesis and organ size control (29, 30, 33). Our work is thus a first step towards more comprehensive models of specific biological systems.

## ACKNOWLEDGMENTS

We thank Alex Golden for helpful conversations. This work was supported by NSF grant DMR-1056456, the Margaret and Herman Sokol Faculty Awards, and the NSF Graduate Research Fellowship Program under grant DGE-1256260.

## Notes

The authors declare no conflict of interest.

