## Supplementary information for "The statistics of noisy growth with mechanical feedback in elastic tissues"

### Supplementary information for:
### The statistics of noisy growth with mechanical feedback in elastic tissues

**Ojan Khatib Damavandi**[*] **and David K. Lubensky**[†]

**Introduction.** In this Supporting Information (SI), we present several derivations and calculations that we skipped in the main text and some theoretical background regarding the quantities of interest. First, we derive our expression for the stress tensor (main text Eq. 1) from the general nonlinear theory of morphoelasticity (1). Next, we show a linearization of the growth dynamics in Eq. 3 leading to Eq. 4 in the main text. Then, we discuss our choice of time derivative and some subtleties associated with material time derivatives of tensors. In the same section, we also discuss the equivalence of our model with the fluidization picture proposed by Ranft et al. (2). After that, we show the full derivation of the spatial density-density correlation function (Eq. 7) and derive an expression for the time-dependent density-density correlation function. We then consider the effect of including gradients of the stress tensor in the growth dynamics and verify our claim in the main text that this is not qualitatively different from the simple stress feedback we presented there. Next, we show the extension of our model for isotropic growth to higher dimensions. Then we move on to anisotropic growth and provide a more detailed derivation of the anisotropic growth dynamics and soft modes (Eqs. 14–16), which we will then use in our calculation of clone statistics. In the final section of this SI, we return to the special case of no net growth, $\gamma_0 = 0$, recovering several results from (2).

Throughout this SI, to avoid confusion, we refer to equations in the main text by their equation number, without any prefix, while for equations in the SI we use 'S' followed by a number. Thus, for example, Equation 1 in the main text will be referred to as Eq. 1, and Equation 1 in the SI will be called Eq. S1.

**Derivation of Cauchy Stress from Nonlinear Morphoelasticity.** In this section, we show how the expression for the Cauchy stress tensor given in Eq. 1 of the main text (and valid in the limit of small deviations from uniform growth) can be obtained by linearizing the general, nonlinear theory of morphoelasticity. We quote here without justification a number of well-established results in morphoelasticity; for derivations and a more in-depth explanation, the interested reader is referred to (1). Following the morphoelasticity literature, we call the fully nonlinear Cauchy stress tensor $T_{ij}$, reserving $\sigma_{ij}$ for the linearized version used in the main text.

The theory of morphoelasticity is an extension of finite strain theory, applied when deformations can no longer be considered infinitesimal. Morphoelasticity deals with arbitrary deformations due to growth. The deformation gradient $F_{ij} = \partial r_i/\partial R_j$ is defined which maps the Eulerian to Lagrangian coordinates. The underlying assumption of morphoelasticity (3) is that we can decompose the deformation gradient into a growth part G followed by an elastic deformation denoted by A, F = AG. Then, the Cauchy stress tensor is taken to be related in the usual manner to the elastic part of the deformation:

$$\mathrm{T} = J^{-1}\mathrm{A}\frac{\partial W}{\partial \mathrm{A}} = 2J^{-1}\mathrm{A}\frac{\partial W}{\partial \mathrm{A}^T\mathrm{A}}\mathrm{A}^T$$

where $J = \det(\mathrm{A})$ and $W$ is the elastic energy density per unit volume of the so-called virtual configuration, which we can imagine as the state of the material after growth but before any elastic deformations. For an isotropic, neo-Hookean material,

$$W = \frac{1}{2}\mathcal{A}_{ijkl}\ \epsilon_{ij}\ \epsilon_{kl} = \frac{1}{2}(\lambda\ \epsilon_{ll}^2 + 2\mu\ \epsilon_{ij}^2),$$

where $\mathcal{A}_{ijkl} = \lambda\delta_{ij}\delta_{kl} + \mu(\delta_{ik}\delta_{jl} + \delta_{il}\delta_{jk})$ is the elasticity tensor for an isotropic body, $\epsilon = (\mathrm{A}^T\mathrm{A} - \mathbb{1})/2$ is the morphoelastic strain tensor, and summation over repeated indices is implied. It is helpful to express A in terms of F and G. Therefore we define a new strain tensor

$$\epsilon' = \mathrm{G}^T\epsilon\ \mathrm{G} = (\mathrm{F}^T\mathrm{F} - \mathrm{G}^T\mathrm{G})/2.$$

Then with the new elastic tensor $\mathcal{A}'_{ijkl} = \lambda(\mathrm{G}^T\mathrm{G})^{-1}_{ij}(\mathrm{G}^T\mathrm{G})^{-1}_{kl} + \mu\left((\mathrm{G}^T\mathrm{G})^{-1}_{ik}(\mathrm{G}^T\mathrm{G})^{-1}_{jl} + (\mathrm{G}^T\mathrm{G})^{-1}_{il}(\mathrm{G}^T\mathrm{G})^{-1}_{jk}\right)$, the energy density $W = \mathcal{A}'_{ijkl}\ \epsilon'_{ij}\ \epsilon'_{kl}/2$.

Now we linearize for small $\widetilde{G}_{ij}/\bar{G}$ (and $w_{ij}/\bar{G}$ because deformations are of the same order as growth fluctuations) to get the results in the limit we are interested in. First, we note that $\mathrm{A} = \mathbb{1} + \mathcal{O}(\widetilde{\mathrm{G}}/\bar{G})$ because uniform growth does not cause any deformations. Then $\epsilon = \mathcal{O}(\widetilde{\mathrm{G}}/\bar{G})$ and we can approximate

$$\mathrm{T} \approx 2\ \partial W/\partial(\mathrm{A}^T\mathrm{A}) = \partial W/\partial\epsilon. \tag{S1}$$

To linear order in $w_{ij}/\bar{G}$ and $\widetilde{G}_{ij}/\bar{G}$, $(\mathrm{F}^T\mathrm{F})_{ij} = \partial_i\mathbf{r}\ .\ \partial_j\mathbf{r} \approx \bar{G}^2\delta_{ij} + 2\bar{G}\ w_{ij}$ and $(\mathrm{G}^T\mathrm{G})_{ij} \approx \bar{G}^2\delta_{ij} + 2\bar{G}\ \widetilde{G}_{ij}$ respectively. Therefore, we can explicitly write out $\epsilon'_{ij}$:

$$\epsilon'_{ij} \approx \bar{G}^2\epsilon_{ij} \approx \bar{G}(w_{ij} - \widetilde{G}_{ij}),$$

[*]
[†]

and so $\epsilon_{ij} \approx (w_{ij} - \widetilde{G}_{ij})/\bar{G}$. This allows us to express $W$ and thus T in terms of $w_{ij}$ and $\widetilde{G}_{ij}$. The elastic energy density to linear order in $w_{ij}/\bar{G}$ and $\widetilde{G}_{ij}/\bar{G}$ is

$$W \approx \frac{1}{2\bar{G}^2}\left[\lambda(w_{ll} - \widetilde{G}_{ll})^2 + 2\mu(w_{ij} - \widetilde{G}_{ij})^2\right].$$

Using Eq. S1, the linearized Cauchy stress tensor is

$$T_{ij} \approx \sigma_{ij} = \frac{1}{\bar{G}}\left[\lambda(w_{ll} - \widetilde{G}_{ll})\delta_{ij} + 2\mu(w_{ij} - \widetilde{G}_{ij})\right], \tag{S2}$$

which is Eq. 1 in the main text.

As a side note, the target metric formalism, an equivalent framework to morphoelasticity, also leads to the same result (4). In fact, the new strain tensor $\epsilon'$ that we defined is the strain tensor used in that formalism. In target metric formalism, a target metric $\bar{g}_{ij}$ is defined denoting the grown, stress free configuration, which often cannot be embedded in real space. $\bar{g}_{ij} = (\mathrm{G}^T\mathrm{G})_{ij}$ in morphoelasticity formalism. The metric defining the final configuration (denoted by $g_{ij}$) is indeed $(\mathrm{F}^T\mathrm{F})_{ij}$, and the strain tensor is $\epsilon'_{ij} = (g_{ij} - \bar{g}_{ij})/2$. We found it easier to work with $\epsilon'_{ij}$ than with $\epsilon_{ij}$ because we had explicit expressions for $g_{ij}$ and $\bar{g}_{ij}$ to linear order.

**Linearized Growth Dynamics.** In this section, we show how to get to the linearized growth dynamics equation (Eq. 4 in the main text) starting from the tensorial dynamics in Eq. 3.

$\Gamma$ in Eq. 3 could be a 4th rank tensor in the most general case, but here we show that it simplifies to a 2nd rank tensor in the limit of small fluctuations. In particular we write:

$$\Gamma_{ijkl} = \gamma_0\delta_{ij}\delta_{kl} + K_{ijkl}(\sigma) + \xi_{ijkl}$$

where $K_{ijkl}(\sigma)$ is first order in the stress tensor and $\xi_{ijkl}$ is the noise. Then, knowing $\partial_t\bar{G} = \gamma_0\bar{G}$ we find from $\partial_t\mathrm{G} = \Gamma\mathrm{G}$

$$\partial_t\widetilde{G}_{ij} = \gamma_0\widetilde{G}_{ij} + K_{ijkl}G_{kl} + \xi_{ijkl}G_{kl}$$

or

$$\partial_t\left[\frac{\widetilde{G}_{ij}}{\bar{G}}\right] = (K_{ijkl} + \xi_{ijkl})\left(\delta_{kl} + \frac{\widetilde{G}_{kl}}{\bar{G}}\right)$$

Since $K_{ijkl}$ is first order in the stress tensor and thus $\mathcal{O}(\widetilde{G}_{ij}/\bar{G})$, we can ignore $K_{ijkl}\widetilde{G}_{kl}/\bar{G}$. Thus, to lowest order in $\widetilde{G}_{kl}/\bar{G}$, the most general form of feedback allowed by symmetries of the system is $K_{ijkl}\delta_{kl} = K_{ijll} \approx c\ \sigma_{ll}\ \delta_{ij}/2 + c^{(d)}\ \sigma^{(d)}_{ij}$, where $\sigma^{(d)}_{ij}$ denotes the traceless part of the stress tensor and the superscript $(d)$ stands for deviatoric. The noise term is evaluated in the weak noise limit, meaning that we can write $G_{kl} \approx \bar{G}\delta_{kl}$ and $\xi_{ijkl}G_{kl}/\bar{G} \approx \xi_{ijkl}\delta_{kl} = \xi_{ij}$ where $\xi_{ij}$ is now a 2nd rank tensorial noise. Putting all of this together we arrive at the following growth equation, Eq. 4 in the main text:

$$\partial_t\left[\frac{\widetilde{G}_{ij}}{\bar{G}}\right] = c\ \sigma_{ll}\ \frac{\delta_{ij}}{2} + c^{(d)}\sigma^{(d)}_{ij} + \xi_{ij}. \tag{S3}$$

**On the Choice of Time Derivative and Connection to Ranft et al. (2).** In this section and this section only, we redefine $\partial_t$ to be the time derivative at fixed Eulerian, not Lagrangian, coordinates. Therefore, the time derivative at fixed Lagrangian coordinates used in every other section becomes $\partial_t + \mathbf{v}.\nabla_\mathbf{r} \equiv \mathrm{d}/\mathrm{d}t$, where $\mathbf{v}$ is the velocity of dilation (i.e. flow velocity).

In specifying the dynamics of the tensor G in the main text, we suggested that it should have the form $\mathrm{dG}/\mathrm{d}t = \Gamma\mathrm{G}$ (compare Eq. 3; here, as just explained, we have written the time derivative at fixed Lagrangian coordinates as $\mathrm{d}/\mathrm{d}t$). It has been argued, however (e.g. ref. (2)), that a better choice would be $\mathrm{DG}/\mathrm{D}t = \Gamma\mathrm{G}$, where $\mathrm{D}/\mathrm{D}t$ is the convected corotational time derivative defined, for any tensor $A_{ij}$, as $(\mathrm{D}A_{ij}/\mathrm{D}t) = \partial_t A_{ij} + v_l\partial_{r_l}A_{ij} + \omega_{il}A_{lj} + \omega_{jl}A_{il}$, where $v_i$ is the velocity and $\omega_{ij} = (\partial_{r_i}v_j - \partial_{r_j}v_i)/2$ is the vorticity of the flow. This derivative differs from $\mathrm{d}/\mathrm{d}t$ by the terms proportional to $\omega_{ij}$. In the most general case, the choice between these time derivatives is a subtle one, which potentially depends both on the biological assumptions one wants to make and on technical questions like whether $G_{ij}$ is defined to include rigid rotations. (See, e.g., ref. (1) for more on the question of rigid rotations and the growth tensor.) To leading order in small deviations from uniform growth (and thus in small displacements), however, it turns out we can sidestep this issue entirely: the two time derivatives agree to this order, as we now demonstrate.

First, notice that any time derivative of G contains a derivative of $\bar{G}$ and a derivative of $\widetilde{G}$. The former is the same for all of the possible time derivatives, because $\bar{G}$ has no space dependence and the parts proportional to $\omega_{ij}$ must vanish because of its antisymmetry. For the part proportional to $\widetilde{G}$, to leading order we can drop any terms where $\widetilde{G}$ is multiplied by something small. Now the velocity field is $v_l = \gamma_0 r_l + \mathcal{O}(w_l)$. Thus, noting that $w_l$ is first order small and that $\gamma_0 r_l$ is an irrotational flow, we conclude that $\omega_{ij}$ is first order small and so can be dropped when it multiplies $\widetilde{G}$. We are then left with the bare time derivative and a convection term proportional to $\gamma_0$ (i.e. $\partial_t + \gamma_0 r_l\partial_{r_l}$), which are the same for the two proposed choices $\mathrm{d}/\mathrm{d}t$ and $\mathrm{D}/\mathrm{D}t$.

Now, we show that, in the limit of small growth non-uniformities, our formalism is equivalent to that of Ranft et al., who show fluidization of growing tissues. In particular, we derive Eqs. 12 and 13 in their paper. Working in Eulerian coordinates, they show in Eq. 12 that in a growing tissue, the trace of the stress tensor follows

$$\frac{\mathrm{d}}{\mathrm{d}t}\sigma_{ll} = 2\chi[v_{ll} - \kappa(\rho)], \quad \text{[S4]}$$

where $\mathrm{d}/\mathrm{d}t = \partial_t + \mathbf{v}.\nabla_{\mathbf{r}}$ is the convected time derivative, $\chi = \lambda + \mu$ is the bulk modulus in two dimensions, $\kappa$ the growth rate, and $v_{ij}$ is the rate of strain tensor, defined with respect to the Eulerian coordinates. This means that in contrast to our spatial derivatives, theirs is taken with respect to $r_i$. In Eq. 13 of their paper, they show that the traceless part of the stress tensor relaxes, which they use to conclude that the tissue acts viscoelastic:

$$\left(1 + \tau_a \frac{\mathrm{D}}{\mathrm{D}t}\right) \sigma_{ij}^{(d)} = 2\tau_a \mu\, v_{ij}^{(d)}, \quad \text{[S5]}$$

where $\tau_a$ is the relaxation time.

First, we simplify their equation for the trace, Eq. S4, in the limit of small growth non-uniformities. They have shown in their Eq. 11 that near the isotropic homeostatic state, the trace of stress relaxes in a similar fashion to the traceless part, with relaxation time $\tau$. Here, we show that we only really need mostly uniform growth with small deviations to see the relaxation.

Looking at Eq. S4, we can see that if $\rho$ is uniform, $\sigma_{ll} = 0$. We use this fact to expand both $v_{ll}$ and $\kappa$ for small non-uniformities (i.e. $\rho = \rho_0 + \delta\rho$ and $\delta\rho \ll \rho_0$) keeping in mind that at $\rho = \rho_0$, $v_{ll} = \kappa(\rho_0) = \bar{\kappa}$. We write $\kappa = \bar{\kappa} + \delta\kappa$ and $v_i = \bar{v}_i + \delta v_i$ with $\bar{v}_i = \gamma_0 r_i$ and $\bar{\kappa} = 2\gamma_0$. From $v_i$ we find $v_{ij} = \gamma_0 \delta_{ij} + \delta v_{ij}$. We also write $\delta\kappa$ in terms of $\delta\rho$ as $\delta\kappa \approx \tau^{-1}\delta\rho/\rho_0$ just as Ranft et al. did in the homeostatic case. We then use $\sigma_{ll} = -2\chi\delta\rho/\rho_0$ from Eq. 6 in their paper to put all of this together and arrive at the following equation that shows relaxation of the trace of stress for almost uniform growth in a similar manner to the traceless part:

$$\left(1 + \tau \frac{\mathrm{d}}{\mathrm{d}t}\right) \sigma_{ll} = 2\tau\chi\delta v_{ll}. \quad \text{[S6]}$$

Now, we bring our attention to our formalism and show that we get the same relaxations as Eqs. S5 and S6. Firstly, we ignore the noise in the growth dynamics and assume the density fluctuations are given as an initial condition. Starting from Eq. S2 (Eq. 1 in the main text), we can take the time derivative to get:

$$\frac{\mathrm{d}}{\mathrm{d}t}\sigma_{ij} = \left[\lambda\left(\frac{\mathrm{d}}{\mathrm{d}t}\left[\frac{w_{ll}}{\bar{G}}\right] - \frac{\mathrm{d}}{\mathrm{d}t}\left[\frac{\widetilde{G}_{ll}}{\bar{G}}\right]\right)\delta_{ij} + 2\mu\left(\frac{\mathrm{d}}{\mathrm{d}t}\left[\frac{w_{ij}}{\bar{G}}\right] - \frac{\mathrm{d}}{\mathrm{d}t}\left[\frac{\widetilde{G}_{ij}}{\bar{G}}\right]\right)\right].$$

$(\mathrm{d}/\mathrm{d}t)[w_{ij}/\bar{G}]$ in our framework is actually $\delta v_{ij}$. The factor of $1/\bar{G}$ comes from the fact that $\delta v_{ij}$ is defined in the Eulerian coordinates and $\partial_{r_i} \approx \partial_{R_i}/\bar{G}$. Replacing $(\mathrm{d}/\mathrm{d}t)[\widetilde{G}/\bar{G}]$ with Eq. S3 and dropping the noise term, we get the following equations for the trace and traceless parts of the stress tensor:

$$\begin{aligned}\left(1 + \frac{1}{2\chi c}\frac{\mathrm{d}}{\mathrm{d}t}\right)\sigma_{ll} &= \frac{1}{c}\delta v_{ll} \\ \left(1 + \frac{1}{2\mu\, c^{(d)}}\frac{\mathrm{d}}{\mathrm{d}t}\right)\sigma_{ij}^{(d)} &= \frac{1}{c^{(d)}}\delta v_{ij}^{(d)}\end{aligned}$$

The first equation is the same as Eq. S6 if we define $\tau = 1/(2\chi c)$. Setting $\tau_a = 1/(2\mu c^{(d)})$, the second equation is also the same as Eq. S5 if we replace $\mathrm{D}/\mathrm{D}t$ with $\mathrm{d}/\mathrm{d}t$ (which we may do to leading order in growth non-uniformities, as explained at the beginning of this section), and notice that $v_{ij}^{(d)} = \delta v_{ij}^{(d)}$, easy to see from $v_{ij} = \gamma_0\delta_{ij} + \delta v_{ij}$.

**Isotropic Density-density Correlations.** In the main text we claimed that the spatial density-density correlation shows a power law behavior (Eq. 7) while the time correlations decay exponentially. Here, we show the full calculations and express the correlation functions including the prefactors, which we omitted in the main text. We will only focus on the isotropic case as the dynamics of density fluctuations is the same in isotropic and anisotropic growth, and effectively we just need to change $k \to k + k^{(d)}$ to go from isotropic to anisotropic, as can be seen by comparing Eqs. 6 and 14.

Starting from Eq. 6, we first do the calculations in fixed Lagrangian coordinates to bypass the complexities associated with large convected terms due to the uniform tissue dilation. $\xi$ is naturally defined in the Eulerian coordinates, but in the weak noise limit, we can easily find the noise correlators in the Lagrangian coordinates. First, let us naïvely take the noise to be delta correlated in both time and space, so that in Eulerian coordinates $\langle\xi(\mathbf{r},t)\xi(\mathbf{r}',t')\rangle = D\delta(\mathbf{r}-\mathbf{r}')\delta(t-t')$. In Lagrangian coordinates, we approximate $\mathbf{r} \approx \bar{G}\mathbf{R}$ and write $\langle\xi(\mathbf{R},t)\xi(\mathbf{R}',t')\rangle = D\delta(\mathbf{R}-\mathbf{R}')\delta(t-t')/\bar{G}^2$, where $D$ is the noise strength. We now solve for $\delta\rho(\mathbf{R},t)$ from Eq. 6:

$$\delta\rho(\mathbf{R},t) = \rho_0\frac{\mu}{\lambda+2\mu}e^{-kt}\int_0^t \xi(\mathbf{R},t_1)e^{kt_1}\mathrm{d}t_1.$$

Here, the initial condition is omitted because it will not matter at long time, where we expect to reach a steady state. Then, for the correlation function, we have:

$$\begin{aligned}\frac{1}{\rho_0^2}\langle\delta\rho(\mathbf{R},t)\delta\rho(\mathbf{R}',t)\rangle &= e^{-2kt}\frac{\mu^2}{(\lambda+2\mu)^2}\int_0^t \mathrm{d}t_1\ \mathrm{d}t_2\left\langle\xi(\mathbf{R},t_1)\xi(\mathbf{R}',t_2)\right\rangle e^{k(t_1+t_2)} \\ &= \frac{\mu^2}{(\lambda+2\mu)^2}De^{-2kt}\int_0^t \mathrm{d}t_1\delta(\mathbf{R}-\mathbf{R}')e^{-2\gamma_0 t_1}e^{2kt_1} \qquad \text{[S7]}\end{aligned}$$

From now on use $D'=\mu^2D/(\lambda+2\mu)^2$. Because we want to find the correlation at fixed Eulerian coordinates at time $t$, we can write $\mathbf{R}=e^{-\gamma_0 t}\mathbf{r}$:

$$\frac{1}{\rho_0^2}\langle\delta\rho(\mathbf{r},t)\delta\rho(\mathbf{r}',t)\rangle=\frac{D'}{2(\gamma_0-k)}\left(e^{2(\gamma_0-k)t}-1\right)\delta(\mathbf{r}-\mathbf{r}').$$

The long time behavior of this expression is pathological: for $k<\gamma_0$, the correlation function blows up as $t\to\infty$ while for $k>\gamma_0$ it tends to a delta function. The reason for this pathological behavior is that the dynamics actually never reaches steady state for fixed Eulerian coordinates due to the fact that early fluctuations that are delta correlated in space never have time to reach any finite distance in Eulerian coordinates even if $t\to\infty$.

Therefore, we need to consider a small correlation length for the noise to regularize the growth as we did in the main text. In Lagrangian coordinates, the noise correlator now becomes $\langle\xi(\mathbf{R},t)\xi(\mathbf{R}',t')\rangle=De^{-\frac{\bar{G}(t)^2(\mathbf{R}-\mathbf{R}')^2}{a^2}}\delta(t-t')/\pi a^2$, where $a$ is a small length scale for correlations. Eq. S7 now yields

$$\frac{1}{\rho_0^2}\langle\delta\rho(\mathbf{R},t)\delta\rho(\mathbf{R}',t)\rangle=\frac{D'}{\pi a^2}\ e^{-2kt}\int_0^t \mathrm{d}t_1 e^{-\frac{(\mathbf{R}-\mathbf{R}')^2}{a^2}e^{2\gamma_0 t_1}}e^{2kt_1},$$

or in Eulerian coordinates

$$\frac{1}{\rho_0^2}\langle\delta\rho(\mathbf{r},t)\delta\rho(\mathbf{r}',t)\rangle=\frac{D'}{\pi a^2}\int_0^t \mathrm{d}t_1 e^{-\frac{(\mathbf{r}-\mathbf{r}')^2}{a^2}e^{-2\gamma_0(t-t_1)}}e^{-2k(t-t_1)}.$$

With the change of variable $u=e^{-2\gamma_0(t-t_1)}$ and setting $r'=0$ without loss of generality, we can see that for $t\to\infty$ (i.e. in steady state), the integral simplifies to $\int_0^1 \mathrm{d}u\, u^{k/\gamma_0-1}e^{-r^2u/a^2}$, which is in fact the integral representation of the lower incomplete gamma function defined as $\gamma(s,x)=\int_0^x u^{s-1}e^{-u}\mathrm{d}u$. The full expression for the density-density correlation function is

$$\frac{1}{\rho_0^2}\langle\delta\rho(\mathbf{r},t)\delta\rho(\mathbf{0},t)\rangle_{t\to\infty}=\frac{D'}{(2\gamma_0)\pi a^2}\ \gamma\left(k/\gamma_0,(r/a)^2\right)\left(\frac{r}{a}\right)^{\frac{-2k}{\gamma_0}}\xrightarrow{r\gg a}\frac{D'}{(2\gamma_0)\pi a^2}\ \Gamma\left(\frac{k}{\gamma_0}\right)\left(\frac{r}{a}\right)^{\frac{-2k}{\gamma_0}}, \qquad \text{[S8]}$$

where we have used the fact that for $r\gg a$, $\gamma\left(k/\gamma_0,(r/a)^2\right)$ tends to $\Gamma\left(k/\gamma_0\right)$. Compare this to Eq. 7 in the main text.

We can also estimate the prefactor. Firstly, we expect the tissue to behave similarly under bulk and shear strains so that $\mu^2/(\lambda+2\mu)^2$ is of order 1 and $D'\sim D$. For the purposes of a first estimate, we assume that cells divide independently according to a Poisson process and that they instantaneously double their size upon division. Each division then contributes a fixed area of order $\pi a^2/2$ to the tissue. It is a standard result (e.g. (5)) that this discretized, Poissonian growth process can be approximated by a Langevin equation with noise strength $D=(2\gamma_0)\pi a^2/8$. We emphasize that this is only a very rough estimate for the prefactor because in reality, divisions are not perfectly random, and cells add mass throughout the cell cycle rather than only at the moment of division. With this in mind, our estimate for the prefactor is

$$\frac{D'}{(2\gamma_0)\pi a^2}\ \Gamma\left(\frac{k}{\gamma_0}\right)\sim\frac{1}{8}\Gamma\left(k/\gamma_0\right).$$

For $k\sim\gamma_0$, $\Gamma\left(k/\gamma_0\right)\sim 1$, while for $k\ll\gamma_0$ or $k\gg\gamma_0$, $\Gamma\left(k/\gamma_0\right)\to\infty$. We note, however, that for the case of $k\gg\gamma_0$ (strong feedback), the expression of Eq. S8 as a whole tends to zero, which is expected from a strong feedback.

Finally, we consider density-density time correlations, i.e., correlation between a point initially at $\mathbf{R}$ and itself at a later time $\tau$. This means that in Eulerian coordinates, we are looking at two different points, i.e. $(\mathbf{r},t)$ and $(\mathbf{r}',t+\tau)$, such that both points originate from $(\mathbf{R},0)$. We show that this correlator decays exponentially in time as expected from the negative feedback. Taking $t'=t+\tau$, we can see that

$$\frac{1}{\rho_0^2}\langle\delta\rho(\mathbf{R},t)\delta\rho(\mathbf{R},t+\tau)\rangle=\frac{D'}{\pi a^2}\ e^{-k\tau}\int_0^t \mathrm{d}t_1 e^{2k(t_1-t)}=\frac{D'}{(2k)\pi a^2}\ e^{-k\tau}\left(1-e^{-2kt}\right).$$

Again, we are interested in long time behavior ($t\gg k^{-1}$), which shows exponential decay, $\langle\delta\rho(\mathbf{R},t)\delta\rho(\mathbf{R},t+\tau)\rangle\sim e^{-k\tau}$; using the same estimate for $D'$ as in the previous paragraph, we find that the prefactor is roughly $\gamma_0/8k$.

**Laplacian Feedback.** In the main text and above, we only considered mechanical feedbacks proportional to stress. However, in principle, spatial derivatives of stress could also feed back on the growth. Continuing to work to linear order in the stress, the lowest order term allowed by symmetry in a gradient expansion is the Laplacian of the stress. We show here that this term has the effect of regularizing the model's short distance behavior so that the correlation functions are well-behaved in the limit that the noise is delta-function correlated in space. In particular, we explicitly calculate the density-density correlation functions for isotropic growth and find that they exhibit the same power law behavior at large distances as we found in the main text without the Laplacian feedback but with noise that is colored in space. Because cells most naturally measure local stress differences in the current state of the tissue, not with respect to the initial state, the Laplacian should be taken with respect to the Eulerian coordinates. Eq. 6 must then be modified to read

$$\partial_t \delta\rho(\mathbf{R},t) = -k_1\delta\rho + k_2\nabla^2_{\mathbf{r}}\delta\rho + \rho_0\frac{\mu}{\lambda+2\mu}\xi(\mathbf{R},t), \qquad \text{[S9]}$$

where $k_2$ is the strength of Laplacian feedback, and the noise is chosen to be delta correlated in time and space, i.e. $\langle\xi(\mathbf{r},t)\xi(\mathbf{r}',t')\rangle = D\ \delta(t-t')\ \delta(\mathbf{r}-\mathbf{r}')$ or in Lagrangian coordinates $\langle\xi(\mathbf{R},t)\xi(\mathbf{R}',t')\rangle = D\ \delta(t-t')\ \delta(\mathbf{R}-\mathbf{R}')/\bar{G}^2$, as before. Note that $k_2>0$ otherwise the dynamics would not be stable. To leading order in $\delta\rho$, we can approximate $\nabla^2_{\mathbf{r}}\delta\rho \approx \nabla^2_{\mathbf{R}}\delta\rho/\bar{G}^2$ because $\delta\rho$ is of the same order as $\mathbf{w}/\bar{G}$.

To solve Eq. S9 we go to Fourier space, using the convention

$$f(\mathbf{Q},t) = \int \mathrm{d}\mathbf{R}\ e^{-i\mathbf{Q}.\mathbf{R}} f(\mathbf{R},t)\ ,$$

and define a particular solution

$$\delta\rho^{(p)}(\mathbf{Q},t) = e^{-k_1 t}\ e^{\frac{k_2 Q^2}{2\gamma_0\bar{G}(t)^2}}\ e^{-\frac{k_2 Q^2}{2\gamma_0}}\,.$$

Then, it is easy to see that the full solution of Eq. S9 in Fourier space is

$$\delta\rho(\mathbf{Q},t) = \rho_0\frac{\mu}{\lambda+2\mu}\ \delta\rho^{(p)}(\mathbf{Q},t)\int_0^t \frac{\xi(\mathbf{Q},t_1)}{\delta\rho^{(p)}(\mathbf{Q},t_1)}\mathrm{d}t_1$$

with $\langle\xi(\mathbf{Q},t)\xi(\mathbf{Q}',t')\rangle = (2\pi)^2 D\ \delta(t-t')\ \delta(\mathbf{Q}+\mathbf{Q}')/\bar{G}^2$. We can then find $\langle\delta\rho(\mathbf{Q},t)\delta\rho(\mathbf{Q}',t)\rangle$. We are interested to find $\langle\delta\rho(\mathbf{r},t)\delta\rho(\mathbf{0},t)\rangle_{t\to\infty}$, but we need to be careful about when to take the limit $t\to\infty$. We proceed as follows: First, we take the inverse Fourier transform to find $\langle\delta\rho(\mathbf{R},t)\delta\rho(\mathbf{R}',t)\rangle$, then make the change $\mathbf{R}=\mathbf{r}/\bar{G}(t)$ and $\mathbf{R}'=\mathbf{r}'/\bar{G}(t)$ to arrive at

$$\frac{1}{\rho_0^2}\langle\delta\rho(\mathbf{r},t)\delta\rho(\mathbf{r}',t)\rangle = \frac{D'}{(2\pi)^2}\int_0^t \frac{\mathrm{d}t_1}{\bar{G}(t_1)^2}e^{-2k_1(t-t_1)}\int \mathrm{d}\mathbf{Q}\ e^{\frac{i\mathbf{Q}.(\mathbf{r}-\mathbf{r}')}{\bar{G}(t)}}\ e^{\frac{k_2}{\gamma_0}\left(\frac{1}{\bar{G}(t)^2}-\frac{1}{\bar{G}(t_1)^2}\right)Q^2}\,.$$

Here, it is easier to take the $Q$ integral first. Also setting $\mathbf{r}'=0$ we find

$$\frac{1}{\rho_0^2}\langle\delta\rho(\mathbf{r},t)\delta\rho(\mathbf{0},t)\rangle = \frac{D'\ \gamma_0}{4\pi k_2}\int_0^t \mathrm{d}t_1\ e^{-2k_1(t-t_1)}\frac{e^{-\frac{r^2}{\frac{4k_2}{\gamma_0}\left(\frac{\bar{G}(t)^2}{\bar{G}(t_1)^2}-1\right)}}}{\bar{G}(t_1)^2\left(\frac{1}{\bar{G}(t_1)^2}-\frac{1}{\bar{G}(t)^2}\right)}.$$

With the change of variable $t_2=t-t_1$, we eliminate any explicit dependence on $t$ in the integrand, allowing us to easily take the limit $t\to\infty$ in the limits of integration. The resulting integral is

$$\frac{1}{\rho_0^2}\langle\delta\rho(\mathbf{r},t)\delta\rho(\mathbf{0},t)\rangle_{t\to\infty} = \frac{D'\ \gamma_0}{4\pi k_2}\int_0^\infty \mathrm{d}t_2\ e^{-2k_1 t_2}\frac{e^{2\gamma_0 t_2}}{e^{2\gamma_0 t_2}-1}\ e^{-\frac{r^2}{\frac{4k_2}{\gamma_0}\left(e^{2\gamma_0 t_2}-1\right)}}.$$

With a final change of variable $y=1/(e^{2\gamma_0 t_2}-1)$, we arrive at

$$\frac{1}{\rho_0^2}\langle\delta\rho(\mathbf{r},t)\delta\rho(\mathbf{0},t)\rangle_{t\to\infty} = \frac{D'}{8\pi k_2}\int_0^\infty \mathrm{d}y\ y^{\frac{k_1}{\gamma_0}-1}(1+y)^{-\frac{k_1}{\gamma_0}}\ e^{-\frac{\gamma_0\ r^2}{4k_2}y},$$

which is the integral representation of the so-called confluent hypergeometric function of the second kind defined as $U(a,b,x) = 1/\Gamma(a)\int_0^\infty \mathrm{d}y\ y^{a-1}(1+y)^{b-a-1}\ e^{-xy}$ with the asymptotic behavior $\lim_{x\to\infty}U(a,b,x)\sim x^{-a}[1+\mathcal{O}(1/x)]$ (6). Defining $a^2 := 4k_2/\gamma_0$, we get

$$\frac{1}{\rho_0^2}\langle\delta\rho(\mathbf{r},t)\delta\rho(\mathbf{0},t)\rangle_{t\to\infty} = \frac{D'}{(2\gamma_0)\pi a^2}\ \Gamma\left(\frac{k_1}{\gamma_0}\right)\ U\left(\frac{k_1}{\gamma_0},1,\left(\frac{r}{a}\right)^2\right). \qquad \text{[S10]}$$

For $r\gg a$, using the asymptotic form of $U(a,b,x)$, we arrive at the same power law as in Eq. S8:

$$\frac{1}{\rho_0^2}\langle\delta\rho(\mathbf{r},t)\delta\rho(\mathbf{0},t)\rangle_{\substack{t\to\infty\\ r\gg a}} = \frac{D'}{(2\gamma_0)\pi a^2}\ \Gamma\left(\frac{k_1}{\gamma_0}\right)\ \left(\frac{r}{a}\right)^{\frac{-2k_1}{\gamma_0}}.$$

As we can see, in this case, the length scale $a = \sqrt{4k_2/\gamma_0}$ was determined by the Laplacian feedback strength instead of by a correlation length for the noise. This length scale could span several cells depending on how strong the feedback is relative to the average growth rate $\gamma_0$. The power law then is regulated purely by $k_1/\gamma_0$ as before. In other words, the role of the Laplacian feedback is to provide a length scale for the early fluctuations to be carried over the tissue as it dilates. We note that the Laplacian feedback with delta correlated noise is only valid for separations greater than cell size: for large distances, we can take the limit of cell size $\to 0$ and use a delta correlated noise, with Laplacian feedback providing a correlation length $\sqrt{4k_2/\gamma_0}$ for the dynamics; however, for distances close to a cell size, we cannot assume a delta correlated noise anymore and need to have the colored noise as before to regularize the correlations. This is evident by noticing that $U(a, b, x)$ blows up like $\log 1/x$ as $x \to 0$ meaning that without a cut off density-density correlations diverge for $r \to 0$ which is nonphysical.

Fig. S1 compares the density-density correlation function with Laplacian feedback and without ($k_2 = 0$). As can be seen, both follow a power law for large $r/a$, but the Laplacian feedback has a slower convergence to the power law. This plot assumes that both cases have the same length scale $a$, while the source of this length scale is very different: for the Laplacian feedback, it is given by the strength of the feedback, whereas for $k_2 = 0$ it comes from having a colored noise in space.

**Generalization of Isotropic Growth to $d$ dimensions.** It is fairly straightforward to generalize the results for density-density correlation to $d$ dimensions. In this section, we will introduce quantities with subscript $d$ (e.g. $k_d$), which are the $d$ dimensional version of quantities that we have defined before; subscript $d$ should not be confused with superscript $(d)$ that stands for deviatoric and is reserved for traceless tensors or scalars associated with such tensors (e.g. $k^{(d)}$).

Here, we show that the power-law behavior derived in the main text (Eq. 7) and in the Laplacian Feedback section above still hold in arbitrary $d$ dimensions. Writing $\widetilde{G}_{ij} = \widetilde{G}\,\delta_{ij}$, we find from Eq. 2 that

$$w_{ll} = \frac{d\lambda + 2\mu}{\lambda + 2\mu}\widetilde{G}.$$

Then from Eq. 5, we have

$$\delta\rho = \rho_0 \left(\frac{2(d-1)\mu}{\lambda + 2\mu}\right) \frac{\widetilde{G}}{\bar{G}}.$$

Eq. 4 in $d$ dimensions looks like

$$\partial_t \left[\frac{\widetilde{G}_{ij}}{\bar{G}}\right] = c\ \sigma_{ll}\ \frac{\delta_{ij}}{d} + c^{(d)}\sigma^{(d)}_{ij} + \xi_{ij}(\mathbf{R}, t),$$

and so for isotropic growth in $d$ dimensions we get

$$d\,\partial_t \left[\frac{\widetilde{G}}{\bar{G}}\right] = c\ \sigma_{ll} + \xi(\mathbf{R}, t),$$

where $\xi(\mathbf{R}, t) = \xi_{ll}(\mathbf{R}, t)$. Therefore, the dynamics of $\delta\rho$ will be

$$\partial_t \delta\rho = -k_d \delta\rho + \rho_0 \frac{2(d-1)\mu}{d(\lambda + 2\mu)}\xi(\mathbf{R}, t), \qquad \text{[S11]}$$

where $k_d = 2(d-1)\mu(d\lambda+2\mu)c/(d\lambda+2d\mu)$. For $d = 2$, we recover Eq. 6. To solve Eq. S11 and find the density-density correlator, we first need to rewrite the noise correlator in $d$ dimension (and remember that we are in the case of no Laplacian feedback and so need to use a colored noise): $\langle \xi(\mathbf{R}, t)\xi(\mathbf{R}', t')\rangle = D_d\, e^{-\frac{\bar{G}(t)^2(\mathbf{R}-\mathbf{R}')^2}{a^2}}\delta(t - t')/(\pi a^2)^{d/2}$. After some straightforward algebra (see Isotropic Density-density Correlation section), we find:

$$\frac{1}{\rho_0^2}\langle \delta\rho(\mathbf{r}, t)\delta\rho(\mathbf{0}, t)\rangle_{t\to\infty} = \frac{D'_d}{(2\gamma_0)(\pi a^2)^{d/2}}\ \gamma\left(k_d/\gamma_0, (r/a)^2\right)\ \left(\frac{r}{a}\right)^{\frac{-2k_d}{\gamma_0}} \xrightarrow{r \gg a} \frac{D'_d}{(2\gamma_0)(\pi a^2)^{d/2}}\ \Gamma\left(\frac{k_d}{\gamma_0}\right)\ \left(\frac{r}{a}\right)^{\frac{-2k_d}{\gamma_0}}, \qquad \text{[S12]}$$

with $D'_d = [2(d-1)\mu/(d\lambda + 2d\mu)]^2\, D_d$. Comparing this with Eq. S8, we see that the density-density correlator shows the same power-law behavior in any dimensions with $d$-dependent exponent and prefactors. To estimate the prefactor in this case, we follow the same argument presented in Isotropic Density-density Correlation section, namely, we estimate the noise to be due to random Poisson divisions each contributing the same $d$ dimensional volume $\Delta V_d$ to the tissue. In particular, $D_d = \Delta V_d(d\gamma_0)/d^2$. $d\gamma_0$ comes from the fact that in a Poisson-like growth, noise is proportional square root of volumetric growth rate, which is precisely $d\gamma_0$. The factor of $1/d^2$ comes from change of variable from volumetric growth to density. $\Delta V_d$ on the other hand is assumed to be the volume of a $d$ dimensional sphere with radius $a/\sqrt{2}$, or $\Omega_d a^d/(2^{d/2}d)$ where $\Omega_d$ is the solid angle in $d$ dimensions. Therefore, our estimate for $D_d$ is $D_d = \gamma_0\Omega_d\, a^d/(d^2\, 2^{d/2})$.

Now, we show that the same power law of Eq. S12 is achieved with Laplacian feedback and delta correlated noise in $d$ dimensions. The differential equation for $\delta\rho$ is

$$\partial_t \delta\rho = -k_{d,1}\delta\rho + k_{d,2}\nabla^2_{\mathbf{r}}\delta\rho + \rho_0 \frac{2(d-1)\mu}{d(\lambda + 2\mu)}\xi(\mathbf{R}, t), \qquad \text{[S13]}$$

and the noise correlator is $\langle \xi(\mathbf{R},t)\xi(\mathbf{R}',t')\rangle = D\ \delta(t-t')\ \delta(\mathbf{R}-\mathbf{R}')/\bar{G}^d$. We follow the exact same steps we did in Laplacian Feedback section to solve Eq. S13. The only difference is that the $\mathbf{Q}$ integrals are now $d$ dimensional. After some algebra, we get

$$\frac{1}{\rho_0^2}\langle \delta\rho(\mathbf{r},t)\delta\rho(\mathbf{0},t)\rangle = \frac{D_d'\ \pi}{(2\pi)^d}\left(\frac{\gamma_0}{k_{d,2}}\right)^{d/2}\int_0^t \mathrm{d}t_1\ e^{-2k_{d,1}(t-t_1)}\ \frac{e^{-\frac{r^2}{\frac{4k_{d,2}}{\gamma_0}\left(\frac{\bar{G}(t)^2}{\bar{G}(t_1)^2}-1\right)}}}{\bar{G}(t_1)^d\left(\frac{1}{\bar{G}(t_1)^2}-\frac{1}{\bar{G}(t)^2}\right)^{d/2}}.$$

Now we eliminate the explicit $t$ dependence with the change of variable $t_2 = t - t_1$ allowing us to take $t\to\infty$, and do another change of variable $y = 1/(e^{2\gamma_0 t_2}-1)$ just like we did before to find

$$\frac{1}{\rho_0^2}\langle \delta\rho(\mathbf{r},t)\delta\rho(\mathbf{0},t)\rangle_{t\to\infty} = \frac{D_d'\ \pi}{(2\pi)^d(2\gamma_0)}\left(\frac{\gamma_0}{k_{d,2}}\right)^{d/2}\int_0^\infty \mathrm{d}y\ y^{\frac{k_{d,1}}{\gamma_0}-1}(1+y)^{\frac{d}{2}-\frac{k_{d,1}}{\gamma_0}-1}\ e^{-\frac{\gamma_0}{4k_{d,2}}\frac{r^2}{y}}.$$

In terms of $U(a,b,x)$, we have

$$\frac{1}{\rho_0^2}\langle \delta\rho(\mathbf{r},t)\delta\rho(\mathbf{0},t)\rangle_{t\to\infty} = \frac{D_d'}{(2\gamma_0)\pi^{d-1}a^d}\ \Gamma\left(\frac{k_{d,1}}{\gamma_0}\right)\ U\left(\frac{k_{d,1}}{\gamma_0},\frac{d}{2},\left(\frac{r}{a}\right)^2\right) \xrightarrow{r\gg a} \frac{D_d'}{(2\gamma_0)\pi^{d-1}a^d}\ \Gamma\left(\frac{k_{d,1}}{\gamma_0}\right)\ \left(\frac{r}{a}\right)^{\frac{-2k_{d,1}}{\gamma_0}}, \quad \text{[S14]}$$

which shows the same power law as Eq. S10. Here $a=\sqrt{4k_{d,2}/\gamma_0}$. This concludes the extension of isotropic growth to $d$ dimensions.

**Anisotropic Growth Equations.** Going back to $d=2$, here we discuss the decomposition we used for $\widetilde{G}_{ij}$ in Fourier space (Eq. 10), and the derivation of the mode structure and growth equations when anisotropic growth is allowed (Eqs. 14–16).

We first find $\mathbf{w}$ in terms of $\widetilde{G}_{ij}$ from Eq. 2 in Fourier space:

$$(\lambda+2\mu)(\mathbf{Q}.\mathbf{w})\mathbf{Q} - \mu(\mathbf{Q}\times\mathbf{Q}\times\mathbf{w}) = -i[\lambda\ \widetilde{G}_{ll}\ \mathbf{Q} + 2\mu\ \mathbf{Q}.\widetilde{\mathrm{G}}], \quad \text{[S15]}$$

where $[\mathbf{Q}.\widetilde{\mathrm{G}}]_i = Q_j\widetilde{G}_{ij}$. To solve this equation, we proposed the following decomposition, Eq. 10:

$$\widetilde{G}_{ij} = \widetilde{G}_{ll}\frac{\delta_{ij}}{2} + (\widetilde{G}_{\parallel}\delta_{ik} - \widetilde{G}_{\perp}\epsilon_{ik})\left[\frac{2Q_kQ_j}{Q^2}-\delta_{kj}\right].$$

Here, we have decomposed $\widetilde{G}_{ij}$ into the trace and nematic components parallel and perpendicular to $\mathbf{Q}$. $\widetilde{G}_{\parallel}$ and $\widetilde{G}_{\perp}$ are $Q$-dependent, and the traceless symmetric tensor made with them as its basis is related to the traceless part of $\widetilde{G}_{ij}$ by a rotation in $Q$-space, i.e.

$$\begin{bmatrix}\widetilde{G}_{\parallel} & \widetilde{G}_{\perp}\\ \widetilde{G}_{\perp} & -\widetilde{G}_{\parallel}\end{bmatrix} = \begin{bmatrix}\cos\beta & -\sin\beta\\ \sin\beta & \cos\beta\end{bmatrix}\begin{bmatrix}\frac{1}{2}(\widetilde{G}_{11}-\widetilde{G}_{22}) & \widetilde{G}_{12}\\ \widetilde{G}_{12} & \frac{1}{2}(\widetilde{G}_{22}-\widetilde{G}_{11})\end{bmatrix}\begin{bmatrix}\cos\beta & \sin\beta\\ -\sin\beta & \cos\beta\end{bmatrix}, \quad \text{[S16]}$$

where $2\beta = \sin^{-1}(-2Q_1Q_2/Q^2)$. This rotation takes any tensor decomposed in this specific way to a basis without explicit $Q$-dependence, which will prove useful later on when deriving the growth dynamics.

The lefthand side of Eq. S15 is already decomposed into a longitudinal term $(\mathbf{Q}.\mathbf{w})\mathbf{Q}$ and a transverse term $\mathbf{Q}\times\mathbf{Q}\times\mathbf{w}$. Therefore, with the aforementioned decomposition of $\widetilde{G}_{ij}$ we can easily find $\mathbf{w}^{\parallel}$ and $\mathbf{w}^{\perp}$ in terms of the growth tensor as we did in Eq. 11.

Next, we can find the stress tensor in terms of the 3 components of $\widetilde{G}_{ij}$ and then write down the dynamics for these components starting from Eq. 4. One can see easily that the strain tensor in terms of the growth factor is

$$w_{ij} = \frac{1}{\lambda+2\mu}\left((\lambda+\mu)\widetilde{G}_{ll} + 2\mu\ \widetilde{G}_{\parallel}\right)\frac{Q_iQ_j}{Q^2} - \widetilde{G}_{\perp}\epsilon_{ik}\left[\frac{2Q_kQ_j}{Q^2}-\delta_{kj}\right].$$

It's immediately clear that the transverse part of $w_{ij}$ is exactly the same as the transverse part of $\widetilde{G}_{ij}$, which means that the stress tensor $\sigma_{ij} = [\lambda(w_{ll}-\widetilde{G}_{ll})\delta_{ij} + 2\mu(w_{ij}-\widetilde{G}_{ij})]/\bar{G}$ is not going to have a transverse component. After some algebra we get

$$\sigma_{ij} = \frac{2\mu(\lambda+\mu)}{(\lambda+2\mu)\bar{G}}\ (\widetilde{G}_{ll}-2\widetilde{G}_{\parallel})\left[\frac{Q_iQ_j}{Q^2}-\delta_{ij}\right].$$

Noticing that

$$\delta\rho = \frac{\rho_0}{\bar{G}}(\widetilde{G}_{ll}-w_{ll}) = \rho_0\frac{\mu}{\lambda+2\mu}\left(\frac{\widetilde{G}_{ll}-2\widetilde{G}_{\parallel}}{\bar{G}}\right),$$

we can now rewrite the stress tensor above in terms of $\delta\rho$ to arrive at the expression given in Eq. 12.

To solve the growth dynamics equation (Eq. 4), we first go to Fourier space and find ODEs for $\widetilde{G}_{ll}$, $\widetilde{G}_{\parallel}$ and $\widetilde{G}_{\perp}$. To do so, it is convenient to rotate the tensors in $Q$-space with angle $\beta$ to go to the basis where there is no explicit $Q$-dependence as we showed above in Eq. S16. Note that the stress can be written as $\sigma_{ij} = \sigma_{ll}\ \delta_{ij}/2 + (\sigma_{\parallel}\delta_{ik} - \sigma_{\perp}\epsilon_{ik})(2Q_kQ_j/Q^2 - \delta_{kj})$ where $\sigma_{\parallel} = \mu(\lambda+\mu)(\widetilde{G}_{ll} - 2\widetilde{G}_{\parallel})/[(\lambda+2\mu)\bar{G}]$ and $\sigma_{\perp} = 0$ as stress has no transverse component. Similarly, we write the noise in the same basis as

$$\xi_{ij} = \xi_{ll}\frac{\delta_{ij}}{2} + (\xi_{\parallel}\delta_{ik} - \xi_{\perp}\epsilon_{ik})\left[\frac{2Q_kQ_j}{Q^2} - \delta_{kj}\right]. \qquad \text{[S17]}$$

To find the correlators of $\xi_{ll}$, $\xi_{\parallel}$ and $\xi_{\perp}$, we first need to find the correlators in real space. Note that since Eq. 4 is a first order perturbation about an isotropic growth, we require the noise to be rotationally invariant. Therefore, if we write the noise in real space as

$$\xi_{ij} = \xi_{ll}\frac{\delta_{ij}}{2} + \xi_1\begin{bmatrix}1 & 0\\ 0 & -1\end{bmatrix} + \xi_2\begin{bmatrix}0 & 1\\ 1 & 0\end{bmatrix},$$

where $\xi_{ll}$, $\xi_1$ and $\xi_2$ are independent random variables, then $\xi_1$ and $\xi_2$ need to have the same variance (which could in general be different from the variance of $\xi_{ll}$). In other words, we have the following correlators for the noise components:

$$\begin{aligned}
\langle\xi_{ll}(\mathbf{r},t)\xi_{nn}(\mathbf{r}',t')\rangle &= D_1\frac{e^{-\frac{(\mathbf{r}-\mathbf{r}')^2}{a^2}}}{\pi a^2}\ \delta(t-t'),\\
\langle\xi_1(\mathbf{r},t)\xi_1(\mathbf{r}',t')\rangle &= D_2\frac{e^{-\frac{(\mathbf{r}-\mathbf{r}')^2}{a^2}}}{\pi a^2}\ \delta(t-t'),\\
\langle\xi_2(\mathbf{r},t)\xi_2(\mathbf{r}',t')\rangle &= D_2\frac{e^{-\frac{(\mathbf{r}-\mathbf{r}')^2}{a^2}}}{\pi a^2}\ \delta(t-t'),
\end{aligned}$$

and all cross correlations are zero. Here, we assumed for simplicity that $a$ is the same for $\xi_{1,2}$ and $\xi_{ll}$, but the correlation lengths for these 3 components could be different in general.

In $Q$-space, the correlators will involve $\delta(\mathbf{Q}+\mathbf{Q}')$, therefore different $Q$'s don't mix. This along with rotational invariance leads to statistical independence of $\xi_{ll}$, $\xi_{\parallel}$ and $\xi_{\perp}$, and we get the following correlators:

$$\begin{aligned}
\langle\xi_{ll}(\mathbf{Q},t)\xi_{nn}(\mathbf{Q}',t')\rangle &= (2\pi)^2\ D_1\frac{e^{-(\frac{a\,Q}{2\bar{G}})^2}}{\bar{G}^2}\ \delta(t-t')\ \delta(\mathbf{Q}+\mathbf{Q}'),\\
\langle\xi_{\parallel}(\mathbf{Q},t)\xi_{\parallel}(\mathbf{Q}',t')\rangle &= (2\pi)^2\ D_2\frac{e^{-(\frac{a\,Q}{2\bar{G}})^2}}{\bar{G}^2}\ \delta(t-t')\ \delta(\mathbf{Q}+\mathbf{Q}'),\\
\langle\xi_{\perp}(\mathbf{Q},t)\xi_{\perp}(\mathbf{Q}',t')\rangle &= (2\pi)^2\ D_2\frac{e^{-(\frac{a\,Q}{2\bar{G}})^2}}{\bar{G}^2}\ \delta(t-t')\ \delta(\mathbf{Q}+\mathbf{Q}'),
\end{aligned} \qquad \text{[S18]}$$

with all the cross correlators zero. Now that we have the noise correlators for $\xi_{\parallel}$ and $\xi_{\perp}$, we can apply the rotation in Eq. S16 to both sides of Eq. 4 and find:

$$\partial_t\ \left(\frac{1}{\bar{G}}\right)\begin{bmatrix}\frac{\widetilde{G}_{ll}}{2}+\widetilde{G}_{\parallel} & \widetilde{G}_{\perp}\\ \widetilde{G}_{\perp} & \frac{\widetilde{G}_{ll}}{2}-\widetilde{G}_{\parallel}\end{bmatrix} = -\frac{k}{2\bar{G}}\begin{bmatrix}\widetilde{G}_{ll}-2\widetilde{G}_{\parallel} & 0\\ 0 & \widetilde{G}_{ll}-2\widetilde{G}_{\parallel}\end{bmatrix} + \frac{k^{(d)}}{2\bar{G}}\begin{bmatrix}\widetilde{G}_{ll}-2\widetilde{G}_{\parallel} & 0\\ 0 & -\widetilde{G}_{ll}+2\widetilde{G}_{\parallel}\end{bmatrix} + \begin{bmatrix}\frac{\xi_{ll}}{2}+\xi_{\parallel} & \xi_{\perp}\\ \xi_{\perp} & \frac{\xi_{ll}}{2}-\xi_{\parallel}\end{bmatrix}$$

where $k = 2\mu(\lambda+\mu)c/(\lambda+2\mu)$ and $k^{(d)} = 2\mu(\lambda+\mu)c^{(d)}/(\lambda+2\mu)$. And, finally, by writing $\widetilde{G}_{ll} - 2\widetilde{G}_{\parallel}$ in terms of $\delta\rho$, we arrive at the differential equations describing the growth:

$$\begin{aligned}
\partial_t\ \delta\rho &= -(k+k^{(d)})\delta\rho + \rho_0\frac{\mu}{\lambda+2\mu}\left(\xi_{ll}-2\xi_{\parallel}\right),\\
\partial_t\left[\frac{\widetilde{G}_{\perp}}{\bar{G}}\right] &= \xi_{\perp},\\
\partial_t\left[\frac{\widetilde{G}_{ll}+2\frac{k}{k^{(d)}}\widetilde{G}_{\parallel}}{\bar{G}}\right] &= \xi_{ll}+2\frac{k}{k^{(d)}}\xi_{\parallel}.
\end{aligned}$$

The bottom two equations describe soft modes with diffusive dynamics. We define the amplitudes of the transverse soft mode $Z_T = \widetilde{G}_{\perp}/\bar{G}$ and the longitudinal soft mode $Z_L = [\widetilde{G}_{ll} + (2k/k^{(d)})\widetilde{G}_{\parallel}]/\bar{G}$. The interpretation of these modes is given in the main text. Finally, in terms of these three amplitudes, $\widetilde{G}_{ij}$ is given by

$$\widetilde{G}_{ij} = \bar{G}\left[\frac{k^{(d)}}{k+k^{(d)}}\left(Z_L + \frac{k(\lambda+2\mu)}{k^{(d)}\mu\,\rho_0}\delta\rho\right)\frac{\delta_{ij}}{2} + \frac{k^{(d)}}{2(k+k^{(d)})}\left(Z_L - \frac{\lambda+2\mu}{\mu\,\rho_0}\delta\rho\right)\left(\frac{2Q_iQ_j}{Q^2}-\delta_{ij}\right) - Z_T\epsilon_{ik}\left(\frac{2Q_kQ_j}{Q^2}-\delta_{kj}\right)\right].$$

**Clone Statistics.** In this section, we derive the results for clone size and shape statistics given, for the general case of anisotropic growth, in the main text Eqs. 18 and 19. First, starting from Eq. 8 of the main text, we derive the variance of the clone size (Eq. 18) and show that this variance scales with clone size when there are growth anisotropies but not in the isotropic limit. Next, we follow the same steps for clone shape starting from Eq. 9 to derive the scaling relation in Eq. 19. Finally, we discuss the correlation, or lack thereof, between the areas of two adjacent clones in our model.

***Clone Size.*** To simplify Eq. 8, note that $\nabla.\mathbf{w} = w_{ll}$. In the previous section, we found $w_{ll}$ in $Q$-space to be

$$w_{ll} = \frac{1}{\lambda + 2\mu}\left((\lambda+\mu)\widetilde{G}_{ll} + 2\mu\ \widetilde{G}_{\parallel}\right).$$

We rewrite this expression in terms of $\delta\rho$ and $Z_L$

$$w_{ll} = \bar{G}\left[\alpha_1\frac{\delta\rho}{\rho_0} + \alpha_2 Z_L\right],$$

where $\alpha_1 = [(\lambda+\mu)k - \mu k^{(d)}]/[\mu(k+k^{(d)})]$ and $\alpha_2 = k^{(d)}/(k+k^{(d)})$. As we can see, this quantity has no explicit $Q$ dependence, so we can formally take the inverse Fourier transform of the scalars $w_{ll}$, $\delta\rho$, and $Z_L$. Although it is not easy to interpret $Z_L$ in real space, it is nevertheless useful for us to work in real space. By expressing $w_{ll}$ in terms of $\delta\rho$ and $Z_L$, now in real space, we can write $\mathrm{Var}(A)$ in Eq. 8 as:

$$\mathrm{Var}(A) = \bar{G}^4\int_{R,R'\le R_c}\left[\frac{\alpha_1^2}{\rho_0^2}\left\langle\delta\rho(\mathbf{R},t)\delta\rho(\mathbf{R}',t)\right\rangle + \frac{2\alpha_1\alpha_2}{\rho_0}\left\langle\delta\rho(\mathbf{R},t)Z_L(\mathbf{R}',t)\right\rangle + \alpha_2^2\left\langle Z_L(\mathbf{R},t)Z_L(\mathbf{R}',t)\right\rangle\right]\mathrm{d}\mathbf{R}\,\mathrm{d}\mathbf{R}'.$$

Note that the cross correlation term is not zero because the noises of $\delta\rho$ and $Z_L$ are correlated (see Eqs. 14 and 16). However, as we will shortly see, the long time behavior of the integral is dominated by the $Z_L$ autocorrelation term and the other two terms are negligible in comparison. Let us look at the three correlators one by one. For $\langle\delta\rho(\mathbf{R},t)\delta\rho(\mathbf{R}',t)\rangle$ we have

$$\begin{aligned}\frac{1}{\rho_0^2}\left\langle\delta\rho(\mathbf{R},t)\delta\rho(\mathbf{R}',t)\right\rangle &= e^{-2(k+k^{(d)})t}\frac{\mu^2}{(\lambda+2\mu)^2}\int_0^t \mathrm{d}t_1\,\mathrm{d}t_2\,e^{(k+k^{(d)})(t_1+t_2)}\left[\langle\xi_{ll}(\mathbf{R},t_1)\xi_{ll}(\mathbf{R}',t_1)\rangle + 4\langle\xi_{\parallel}(\mathbf{R},t_1)\xi_{\parallel}(\mathbf{R}',t_1)\rangle\right]\\ &= e^{-2(k+k^{(d)})t}\frac{\mu^2}{(\lambda+2\mu)^2}\frac{D_1+4D_2}{\pi a^2}\int_0^t e^{-\frac{(\mathbf{R}-\mathbf{R}')^2\bar{G}(t_1)}{a^2}}e^{2(k+k^{(d)})t_1}\mathrm{d}t_1.\end{aligned}$$

Notice that here we are working in fixed Lagrangian coordinates because we are only interested in points within a circular clone of Lagrangian radius $R_c$ independent of time. This is in contrast to our calculation of $\langle\delta\rho(\mathbf{r},t)\delta\rho(\mathbf{r}',t)\rangle$ that needed to be carried out in Eulerian coordinates. One benefit of working in Lagrangian coordinates is that we can take the noise to be delta correlated in $\mathbf{r}$ (i.e. work in the $a\to 0$ limit) without introducing any pathological behavior. This simplifies the calculations, so we will take noise to be white in time and space similar to our calculation for Laplacian feedback, i.e.

$$\begin{aligned}\langle\xi_{ll}(\mathbf{R},t)\xi_{nn}(\mathbf{R}',t')\rangle &= \frac{D_1}{\bar{G}^2}\,\delta(t-t')\,\delta(\mathbf{R}-\mathbf{R}'),\\ \langle\xi_{\parallel}(\mathbf{R},t)\xi_{\parallel}(\mathbf{R}',t')\rangle &= \frac{D_2}{\bar{G}^2}\,\delta(t-t')\,\delta(\mathbf{R}-\mathbf{R}'),\\ \langle\xi_{\perp}(\mathbf{R},t)\xi_{\perp}(\mathbf{R}',t')\rangle &= \frac{D_2}{\bar{G}^2}\,\delta(t-t')\,\delta(\mathbf{R}-\mathbf{R}').\end{aligned} \qquad \text{[S19]}$$

With this, we have

$$\frac{1}{\rho_0^2}\left\langle\delta\rho(\mathbf{R},t)\delta\rho(\mathbf{R}',t)\right\rangle = \frac{\mu^2}{(\lambda+2\mu)^2}\,\frac{D_1+4D_2}{2(k+k^{(d)}-\gamma_0)}\left(e^{-2\gamma_0 t} - e^{-2(k+k^{(d)})t}\right)\delta(\mathbf{R}-\mathbf{R}').$$

Similarly, for $\langle\delta\rho(\mathbf{R},t)Z_L(\mathbf{R}',t)\rangle$ we get

$$\frac{1}{\rho_0}\left\langle\delta\rho(\mathbf{R},t)Z_L(\mathbf{R}',t)\right\rangle = \frac{\mu}{\lambda+2\mu}\,\frac{D_1 - 4(k/k^{(d)})D_2}{k+k^{(d)}-2\gamma_0}\left(e^{-2\gamma_0 t} - e^{-(k+k^{(d)})t}\right)\delta(\mathbf{R}-\mathbf{R}').$$

Finally for $\langle Z_L(\mathbf{R},t)Z_L(\mathbf{R}',t)\rangle$ we find

$$\left\langle Z_L(\mathbf{R},t)Z_L(\mathbf{R}',t)\right\rangle = \frac{D_1 + 4(k/k^{(d)})^2 D_2}{2\gamma_0}\left(1 - e^{-2\gamma_0 t}\right)\delta(\mathbf{R}-\mathbf{R}').$$

Before we go any further, we note that $\langle Z_L(\mathbf{R},t)Z_L(\mathbf{R}',t)\rangle$ does not grow linearly with time as one would naïvely expect given the apparently diffusive dynamics of $Z_L$. The reason is that, as Eq. S19 makes clear, the amplitude of the noise decays with time like $1/\bar{G}^2$. This behavior, in turn, arises from the fact that the noise is defined to have constant strength in Eulerian coordinates. But a region of fixed size in Lagrangian coordinates will grow larger and larger in Eulerian coordinates as time progresses, leading overall variation in that region to go down. The growth hence puts a limit on how large the soft mode variances can get, and at long times, $\langle Z_L(\mathbf{R},t)Z_L(\mathbf{R}',t)\rangle$ approaches a constant. (There is in fact one further subtlety here: The noise correlators of the form $\langle \xi(\mathbf{R},t)\xi(\mathbf{R}',t')\rangle$ have an explicit prefactor of $1/\bar{G}^2$ only when we take the limit $a\to 0$ and convert Gaussian correlation functions in space into delta functions. In fact, for finite $a$, the mean-squared mode displacement at a single point $\mathbf{R}$, $\left\langle Z_L(\mathbf{R},t)^2\right\rangle$, does grow linearly in time. The integral of $\langle Z_L(\mathbf{R},t)Z_L(\mathbf{R}',t)\rangle$ over any fixed region in $\mathbf{R}$, however, still approaches a constant, just as in the limit $a\to 0$, because the range of spatial correlations in $\mathbf{R}$ decreases exponentially in time. Similarly, the noise correlations in $\mathbf{Q}$ space, Eq. S18, always have an explicit prefactor of $1/\bar{G}^2$, even for finite $a$, so that the mean-squared value of of $Z_L(\mathbf{Q},t)$ remains bounded for all time.)

Now, if we compare the three correlators at long times, we see that $\langle Z_L(\mathbf{R},t)Z_L(\mathbf{R}',t)\rangle$ dominates the growth as the other two correlators decay exponentially and so can be ignored in the long time limit. In particular, with $\langle A\rangle = \bar{G}^2\pi R_c^2$, as $t\to\infty$ we can see that the ratio $\mathrm{Var}(A)/\langle A\rangle^2$ is a constant:

$$\left.\frac{\mathrm{Var}(A)}{\langle A\rangle^2}\right|_{t\to\infty} = \alpha_2^2\frac{D_1+4(k/k^{(d)})^2D_2}{(2\gamma_0)\pi R_c^2}\sim\frac{1}{R_c^2},$$

agreeing with Eq. 18. Estimating $R_c$ to be of order of a cell radius, the ratio will be of order 1 with the same assumptions on noise strengths as above in the Isotropic Density-density Correlations section. As a final note, this scaling of variance with clone size is the direct result of having the soft mode $Z_L$ and would not occur in isotropic growth. In that case, the only term we have is $\langle\delta\rho(\mathbf{R},t)\delta\rho(\mathbf{R}',t)\rangle$, which does not scale with size, and the ratio will decay to zero at long times.

***Clone Shape.*** Now we move on to the expression for the amplitudes $B_n$ of the modes specifying clone shape, which involves a few subtleties. For one, notice that the integral involves correlators of $w_k$, which was found in $Q$ space and has explicit $Q$ dependence. Therefore, we first need to find the correlator $\langle w_k w_j'\rangle$ in Fourier space, then take the inverse transform and finally take the $\Theta$ integrals.

We first write out the components of $\mathbf{w}$ (Eq. 11) in terms of the modes as we did with $w_{ll}$ for clone size:

$$w_k(\mathbf{Q},t) = -\frac{i\,\bar{G}}{Q^2}\left[(\alpha_1\frac{\delta\rho}{\rho_0}+\alpha_2 Z_L)Q_k - 2Z_T\;\epsilon_{kl}\,Q_l\right].$$

The correlator will look like

$$\begin{aligned}\langle w_k(\mathbf{Q},t)w_j(\mathbf{Q}',t)\rangle &= -\frac{\bar{G}^2}{Q^2\;Q'^2}\Bigg[\left(\frac{\alpha_1^2}{\rho_0^2}\left\langle\delta\rho(\mathbf{Q},t)\delta\rho(\mathbf{Q}',t)\right\rangle+\frac{2\alpha_1\alpha_2}{\rho_0}\left\langle\delta\rho(\mathbf{Q},t)Z_L(\mathbf{Q}',t)\right\rangle+\alpha_2^2\left\langle Z_L(\mathbf{Q},t)Z_L(\mathbf{Q}',t)\right\rangle\right)Q_k\,Q_j'\\ &\quad +4\left\langle Z_T(\mathbf{Q},t)Z_T(\mathbf{Q}',t)\right\rangle\epsilon_{kl}\,\epsilon_{js}Q_l\,Q_s'\Bigg].\end{aligned}$$

Note that $\delta\rho$ and $Z_L$ do not mix with $Z_T$ because their respective noises are statistically independent. We use the correlators in Eq. S19 for noise terms, of which we then take the Fourier transform to go to $Q$-space. The only dominant terms in $\langle w_k(\mathbf{Q},t)w_j(\mathbf{Q}',t)\rangle$ are the soft mode autocorrelations, with the other two terms involving $\delta\rho$ showing the same exponential decay in time as in the clone size calculation above. Therefore we only focus on the two soft mode autocorrelators:

$$\begin{aligned}\left\langle Z_L(\mathbf{Q},t)Z_L(\mathbf{Q}',t)\right\rangle &= (2\pi)^2\frac{D_1+4(k/k^{(d)})^2D_2}{2\gamma_0}\left(1-e^{-2\gamma_0 t}\right)\delta(\mathbf{Q}+\mathbf{Q}'),\\ \left\langle Z_T(\mathbf{Q},t)Z_T(\mathbf{Q}',t)\right\rangle &= (2\pi)^2\frac{D_2}{2\gamma_0}\left(1-e^{-2\gamma_0 t}\right)\delta(\mathbf{Q}+\mathbf{Q}'),\end{aligned}$$

and

$$\langle w_k(\mathbf{Q},t)w_j(\mathbf{Q}',t)\rangle_{t\to\infty}\approx-(2\pi)^2\frac{\bar{G}^2}{Q^4}\left[\alpha_2^2\frac{D_1+4(k/k^{(d)})^2D_2}{2\gamma_0}\;Q_k\,Q_j'+\frac{2D_2}{\gamma_0}\;\epsilon_{kl}\,\epsilon_{js}Q_l\,Q_s'\right]\left(1-e^{-2\gamma_0 t}\right)\delta(\mathbf{Q}+\mathbf{Q}').$$

Now we need to take the inverse Fourier transform before evaluating the integral in Eq. 9. We are basically taking the inverse transform of something like $Q_kQ_j/Q^4$:

$$\begin{aligned}\frac{1}{(2\pi)^4}\int \mathrm{d}\mathbf{Q}\,\mathrm{d}\mathbf{Q}'\,\frac{Q_k\,Q_j'}{Q^4}e^{i(\mathbf{Q}.\mathbf{R}+\mathbf{Q}'.\mathbf{R}')}\delta(\mathbf{Q}+\mathbf{Q}') &= -\frac{1}{(2\pi)^4}\int\mathrm{d}\mathbf{Q}\,\frac{Q_k\,Q_j}{Q^4}e^{i\mathbf{Q}.(\mathbf{R}-\mathbf{R}')}\\ &= \frac{\partial_{R_k}\partial_{R_j}}{(2\pi)^4}\int\mathrm{d}\mathbf{Q}\,\frac{e^{i\mathbf{Q}.(\mathbf{R}-\mathbf{R}')}}{Q^4}.\end{aligned}$$

The inverse transform of $1/Q^4$ is known from the theory of generalized functions (7) to be

$$\int\mathrm{d}\mathbf{Q}\,\frac{e^{i\mathbf{Q}.(\mathbf{R}-\mathbf{R}')}}{Q^4} = -\frac{\pi}{2}\left\{(1-\gamma+\log 2)(\mathbf{R}-\mathbf{R}')^2-(\mathbf{R}-\mathbf{R}')^2\log|\mathbf{R}-\mathbf{R}'|\right\},$$

where $\gamma$ is the Euler's constant and not to be confused with the growth rate $\gamma_0$. The $\log R$ term is allowed by dimensional analysis and turns out to be part of the solution. There is an ambiguity in the scale $l$ in $\log(R/l)$ but it will not affect the final result.

Taking the double derivative of the integral, we get

$$I_{kj} = \partial_{R_k}\partial_{R_j}\int \mathrm{d}\mathbf{Q}\,\frac{e^{i\mathbf{Q}.(\mathbf{R}-\mathbf{R}')}}{Q^4} = \pi\big\{(\gamma - \frac{1}{2} - \log 2)\delta_{kj} + \frac{(R_j - R_j')(R_k - R_k')}{(\mathbf{R}-\mathbf{R}')^2} + \log|\mathbf{R}-\mathbf{R}'|\delta_{kj}\big\}.$$

Now, we just need to put these pieces together to find

$$\langle w_k(\mathbf{R},t) w_j(\mathbf{R}',t)\rangle_{t\to\infty} \approx -\frac{\bar{G}^2}{4\pi\gamma_0}\left[\alpha_2^2 \frac{D_1 + 4(k/k^{(d)})^2 D_2}{2}\; I_{kj} + 2D_2\;\epsilon_{kl}\,\epsilon_{js}\,I_{ls}\right]\left(1 - e^{-2\gamma_0 t}\right),$$

which we then plug into

$$\left\langle |B_n|^2\right\rangle = \frac{1}{(2\pi)^2}\int_0^{2\pi} \mathrm{d}\Theta \mathrm{d}\Theta' \hat{R}_k(\Theta)\hat{R}_j(\Theta')\langle w_k w_j'\rangle e^{-in(\Theta-\Theta')}.$$

The only nonzero term in this integral is given by the $\log|\mathbf{R}-\mathbf{R}'|$ term in $I_{kj}$ and $I_{ls}$. The integral becomes

$$\left\langle |B_n|^2\right\rangle_{t\to\infty} \approx -\frac{\bar{G}^2}{(4\pi)^2\gamma_0}\left[\alpha_2^2\frac{D_1 + 4(k/k^{(d)})^2 D_2}{2} + 2D_2\right]\left(1 - e^{-2\gamma_0 t}\right)\int_0^{2\pi} \mathrm{d}\Theta\,\mathrm{d}\Theta'(\cos\Theta\cos\Theta' + \sin\Theta\sin\Theta')\log|\mathbf{R}-\mathbf{R}'| e^{-in(\Theta-\Theta')}.$$

The final integral was evaluated to be $-2n\pi^2/(n^2-1)$ (for $n > 1$). With $B_0 = \bar{G}R_c$, we can see that in the limit of $t \to \infty$ we get

$$\left.\frac{\left\langle |B_n|^2\right\rangle}{B_0^2}\right|_{t\to\infty} = \left[\alpha_2^2\frac{D_1 + 4(k/k^{(d)})^2 D_2}{4\gamma_0\,R_c^2} + \frac{D_2}{\gamma_0\,R_c^2}\right]\frac{n}{4(n^2-1)} \sim \frac{1}{R_c^2}\frac{n}{n^2-1},$$

In agreement with Eq. 19. Again, assuming $R_c$ is approximately of order of a cell size, the quantity in square brackets will be of order 1. Similar to the clone size, this scaling relation is purely a result of the soft modes. Here, in contrast to the clone size variance, the transverse soft mode is also involved. This is because $w_k$ , which was the important variable here, depends on both soft modes whereas in the case of clone size the important variable is $w_{ll}$ which depends on only the longitudinal soft mode.

***Independence of Adjacent Clone Areas.*** In this subsection, we discuss the correlation between adjacent clones. We claimed in the main text at the end of the Isotropic Growth section that the areas of adjacent clones are uncorrelated in our model. This may sound counterintuitive, especially knowing that due to soft modes, boundaries of clones are soft, which could lead us to expect that one clone can grow at the expense of an adjacent clone. We show here that this in fact is not the case.

To understand the statistical independence of clone areas, it is useful first to consider the simpler situation where we have only a delta-like instantaneous growth at the origin with strength $\nu$, i.e. $G_{ij}(\mathbf{R},0) = (1 + \nu\,\delta(\mathbf{R}))\delta_{ij}$. (Here, $\bar{G} = 1$.) From Eq. 2 in the main text, we can find

$$\mathbf{w}(\mathbf{R},0) = \frac{2\nu(\lambda+\mu)}{\lambda+2\mu}\frac{\mathbf{R}}{R^2},$$

which implies a purely deviatoric strain field except at the origin (8) and thus a localized change in density

$$\delta\rho(\mathbf{R},0) = \rho_0[\widetilde{G}_{ll}(\mathbf{R},0) - w_{ll}(\mathbf{R},0)] = \frac{2\mu\rho_0}{\lambda+2\mu}\nu\delta(\mathbf{R}).$$

This bump in the density induces mechanical feedback, leading to density relaxation. More precisely, the dynamics given by Eqs. 14–16 in the main text but without noise ($\xi_{ij} = 0$), together with Eq. 13 and the initial conditions $\widetilde{G}_{ll}(\mathbf{R},0) = 2\nu\delta(\mathbf{R})$, $\widetilde{G}_{\parallel}(\mathbf{R},0) = \widetilde{G}_{\perp}(\mathbf{R},0) = 0$ allow us to find the growth tensor and the density at time $t$ (note that because there is no explicit Q dependence in Eqs. 14–16, we can formally take the inverse Fourier transform and work in real space):

$$\begin{aligned}
\widetilde{G}_{ll}(\mathbf{R},t) &= \frac{k^{(d)}}{k+k^{(d)}}2\nu\left(1 + \frac{k}{k^{(d)}}e^{-(k+k^{(d)})t}\right)\delta(\mathbf{R})\\
\widetilde{G}_{\parallel}(\mathbf{R},t) &= \frac{k^{(d)}}{k+k^{(d)}}\nu\left(1 - e^{-(k+k^{(d)})t}\right)\delta(\mathbf{R})\\
\widetilde{G}_{\perp}(\mathbf{R},t) &= 0\\
\delta\rho(\mathbf{R},t) &= \frac{2\mu\rho_0}{\lambda+2\mu}\nu e^{-(k+k^{(d)})t}\,\delta(\mathbf{R}).
\end{aligned}$$

Now, from Eq. 11, we can easily see that $\mathbf{w}(\mathbf{R},t) \sim \mathbf{R}/R^2$ stays divergence free ($\nabla.\mathbf{w} \sim \delta(\mathbf{R})$). Therefore, since $\Delta A = \int \nabla.\mathbf{w}\,\mathrm{d}\mathbf{R}$, any region of the tissue that does not contain the origin will not see any increase in area. In other words, if we have two adjacent clones and introduce a small amount of incremental growth at the origin, only the area of the clone containing the origin will increase; the size of the other will be unchanged, though its shape will be distorted (see Fig. 2A in the main text). This implies that the areas of any two clones are uncorrelated. There is an additional subtlety worth mentioning:

While $\widetilde{G}_{ll}(\mathbf{R},t)$ and $\widetilde{G}_{\parallel}(\mathbf{R},t)$ remain local, $\widetilde{G}_{ij}(\mathbf{R},t)$ in general is not localized to the origin. This is due to the fact that $\widetilde{G}_{\parallel}$ lives in Fourier space and does not have a well-defined physical meaning in real space. To see the non-locality of $\widetilde{G}_{ij}(\mathbf{R},t)$, we start from Eq. 10, noticing that $\widetilde{G}_{\parallel}$ is flat in Fourier space and thus, $\widetilde{G}_{ij}(\mathbf{Q},t)$ is $Q$-dependent and not local in real space. For instance, $\widetilde{G}_{12}(\mathbf{Q},t) \propto Q_1Q_2/Q^2$ which yields $\widetilde{G}_{12}(\mathbf{R},t) \propto R_1R_2/R^4$. Nonetheless, since $\nabla.\mathbf{w}$ is localized to the origin (at least in the absence of effects from boundary conditions that we neglect throughout this paper), clone areas remain uncorrelated.

Returning to our full calculation with arbitrary growth tensor G, we can explicitly see this decoupling if we take two adjacent clones of sizes $A_1$ and $A_2$ and look at $\langle \Delta A_1 \Delta A_2 \rangle$. We define $\Delta A_k = \bar{G} \int_{\mathbf{R}_k \in \text{clone } k} \nabla_k.\mathbf{w}_k \, d\mathbf{R}_k$ with $k = 1, 2$, $\mathbf{w}_k = \mathbf{w}(\mathbf{R}_k)$ and $\nabla_k = \nabla_{\mathbf{R}_k}$. Then, the correlation of clone 1 and 2 will be

$$\langle \Delta A_1 \Delta A_2 \rangle = \bar{G}^2 \int \langle \nabla_1.\mathbf{w}_1 \; \nabla_2.\mathbf{w}_2 \rangle \; d\mathbf{R_1} d\mathbf{R_2}$$

This quantity involves noise correlators $\langle \xi_{ll}(\mathbf{R}_1,t)\xi_{ll}(\mathbf{R}_2,t)\rangle$ and $\langle \xi_{\parallel}(\mathbf{R}_1,t)\xi_{\parallel}(\mathbf{R}_2,t)\rangle$ that give $\delta(\mathbf{R}_1 - \mathbf{R}_2)$, and because we are integrating over two separate regions, the integral is zero as claimed in the main text. We note that if instead of delta correlated noise, we consider colored noise with a small width $a$, within our continuum model clones that actually share a boundary must show small but nonzero area correlation because the correlations in the noise stretch across the boundary: $\langle \Delta A_1 \Delta A_2 \rangle \sim \mathcal{O}(L^2 a^2)$ where $L$ is the length of the shared boundary. However, in reality, the interface is where the cells of clone 1 meet the cells of clone 2, and assuming independent divisions of discrete cells, there is no correlation between noise in clone 1 and 2 and $\langle \Delta A_1 \Delta A_2 \rangle$ will again be zero.

**The Limit of No Net Growth.** Here, we derive expressions for fluctuations in density and velocity in the limit of no net growth ($\gamma_0 = 0$ or $\bar{G} = 1$) and show that in this limit, our model is equivalent to the fluctuating homeostatic tissue described in Ranft et al. (2). In particular, we will derive expressions of the same form as Eqs. 21 and 22 in (2), which related the Fourier transformed (in space and time) density and velocity fluctuations $\delta\rho(\mathbf{q},\omega)$ and $\mathbf{v}(\mathbf{q},\omega)$ to appropriate noise terms.

Since we are interested in fluctuations about the steady state of no growth, the distinction between Lagrangian and Eulerian coordinates vanishes to linear order in small quantities, so we will use lower-case $\mathbf{q}$ to denote the wavevector for consistency with (2).

Starting from Eq. 14 of the main text and Fourier transforming in time, we get

$$\left( -i\omega + \frac{2\mu(\lambda+\mu)}{\lambda+2\mu}(c + c^{(d)}) \right) \delta\rho = \rho_0 \frac{\mu}{\lambda+2\mu}(\xi_{ll} - 2\xi_{\parallel}).$$

Now, writing $\lambda + \mu = \chi$, $1/c = 2\chi\tau$ and $1/c^{(d)} = 2\mu\tau_a$, and also noting that the traceless part of the noise tensor $\xi^{(d)}_{ij} = \xi_{ij} - \xi_{ll}\delta_{ij}/2$ is related to $\xi_{\parallel}$ via $\xi_{\parallel} = q_i \xi^{(d)}_{ij} q_j / q^2$ (Eq. S17), we arrive at an expression of the same form as Eq. 21 in (2):

$$\delta\rho = \frac{\tau\,\rho_0(\tau_a\mu)}{(1-i\omega\tau_a)\tau\chi + (1-i\omega\tau)\tau_a\mu} \left[ \xi_{ll} - 2\frac{q_i\xi^{(d)}_{ij}q_j}{q^2} \right]. \qquad \text{[S20]}$$

We note that the extra factors of 4/3 in (2) appear because their calculation was carried out in 3d. The prefactor to $\xi^{(d)}_{ij}$ is also different here from (2) because we have defined the noise to be acting on the growth tensor (see Eq. S3), whereas Ranft et al. have the traceless noise act directly on the traceless stress tensor.

We now find the velocity fluctuations. We have

$$v_k(\mathbf{q},t) = \partial_t w_k(\mathbf{q},t) = -\frac{i}{q^2}\left[ (\alpha_1\,\partial_t \frac{\delta\rho}{\rho_0} + \alpha_2\,\partial_t Z_L) q_k - 2\,\partial_t Z_T\;\epsilon_{kl}\,q_l \right].$$

Denoting the component of velocity parallel to $\mathbf{q}$ as $v^{\parallel} = v_l q_l / q$, we find

$$iq\,v^{\parallel} = \alpha_1 \left[ -\frac{2\mu\chi}{\chi+\mu}\left(\frac{1}{2\tau\chi} + \frac{1}{2\tau_a\mu}\right)\frac{\delta\rho}{\rho_0} + \frac{\mu}{\chi+\mu}\left(\xi_{ll} - 2\frac{q_i\xi^{(d)}_{ij}q_j}{q^2}\right)\right] + \alpha_2\left(\xi_{ll} + 2\frac{\tau_a\mu}{\tau\chi}\,\frac{q_i\xi^{(d)}_{ij}q_j}{q^2}\right),$$

where, in terms of $\tau$ and $\tau_a$, $\alpha_1 = (\tau_a - \tau)\chi/(\tau\chi + \tau_a\mu)$ and $\alpha_2 = \tau\chi/(\tau\chi + \tau_a\mu)$. Using the expression for $\delta\rho$ in Eq. S20 and after some manipulation, we arrive at the following (compare with $v_{\parallel}$ in Eq. 22 of (2))

$$iq\,v^{\parallel} = \frac{1}{(1-i\omega\tau_a)\tau\chi + (1-i\omega\tau)\tau_a\mu}\left[ \tau\chi(1-i\omega\tau_a)\xi_{ll} + \tau_a\mu(1-i\omega\tau)2\frac{q_i\xi^{(d)}_{ij}q_j}{q^2} \right]. \qquad \text{[S21]}$$

Finally, we have, for the component of velocity perpendicular to $\mathbf{q}$

$$v^{\perp}_k = \partial_t w^{\perp}_k = 2i\epsilon_{kl}\frac{q_l}{q^2}\xi_{\perp}.$$

It is easy to see that $\xi_\perp$ is related to $\xi^{(d)}_{ij}$ by $\epsilon_{kl} q_l \xi_\perp = q_k q_m \xi^{(d)}_{mj} q_j / q^2 - \xi^{(d)}_{kj} q_j$. Plugging this in, we obtain

$$v^\perp_k = \frac{2i}{q^2} (q_k q_m \xi^{(d)}_{mj} q_j / q^2 - \xi^{(d)}_{kj} q_j). \tag{S22}$$

Comparing S22 to Eq. 22 in (2), we see that they again only differ by prefactors that can be absorbed in the noise strength by redefinition of $\xi_\perp$.

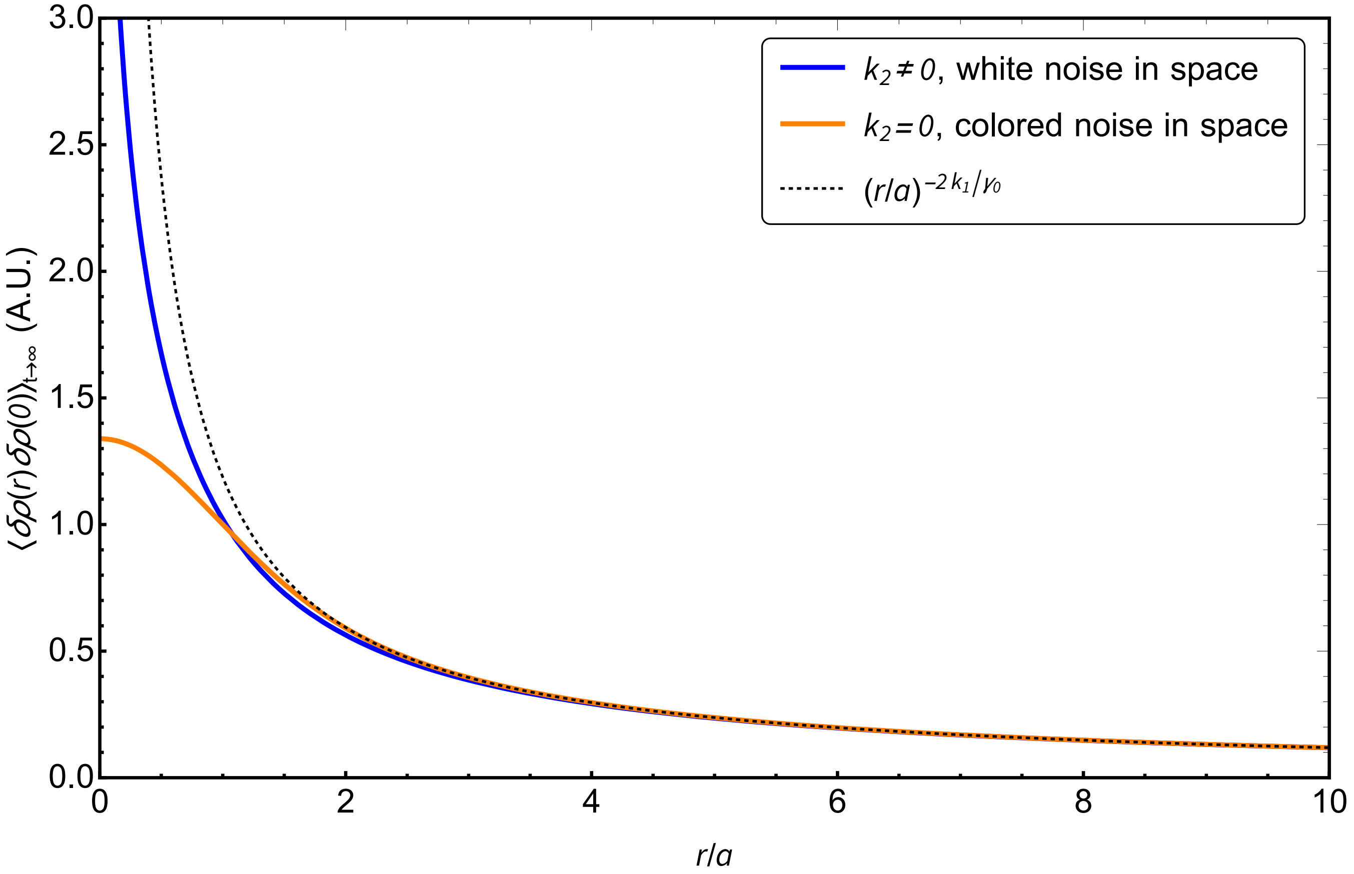


**Fig. S1.** Plot of density-density correlation function for Laplacian feedback ($k_2 \neq 0$) and the simple stress feedback ($k_2 = 0$) discussed in the main text. In the case of $k_2 \neq 0$, the approach to the power law is slower. The plot is for $k_1 = \gamma_0/2$. For $k_2 = 0$, $a$ is the width of the colored noise, whereas for $k_2 \neq 0$, $a = \sqrt{4k_2/\gamma_0}$.
